# Coordinated remodeling of NAD⁺ metabolic networks across the lifespan of *Caenorhabditis elegans*

**DOI:** 10.64898/2026.11.16.767138

**Authors:** Abdulkareem A. Alshaheeb, Mahmoud S. Yahia, Ian A. Argento, Jaedon J. Sadler, Andrew M. Spence, Shaina V. Johnson, Emily R. Ritter, Isabella J. Munoz, Kayla N. Dennis, Brett C. Breidor, Emmanuel Olayiwola, Tatiana N. Laremore, Melanie R. McReynolds

## Abstract

Changes in the metabolic network that sustain NAD^+^ during aging remain poorly understood. Analysis of steady state levels is insufficient to reveal whether biosynthetic or consumption pathways are perturbed or preserved. Here, we address this gap in understanding by integrating metabolomics, proteomics, gene-expression profiling, and stable-isotope tracing across the *Caenorhabditis elegans* adult lifespan. NAD⁺ abundance increases with age despite reductions in selected biosynthetic components. Tracing reveals reduced mid-adult incorporation of nicotinamide and nicotinic acid into NAD⁺, alongside declining nicotinamide riboside abundance and loss of detectable nicotinic acid-derived labeling into this precursor. Early mitochondrial Complex I remodeling accompanies sustained changes in redox and energetic indices. Disruption of NAD⁺-consuming or salvage enzymes reduces reproduction and lifespan, whereas precursor supplementation provides no longevity benefit under the conditions tested. Together, these findings reveal that NAD⁺ accumulation can coexist with extensive metabolic remodeling, establishing the importance of evaluating network dynamics beyond cofactor abundance.

**Highlights:**

- NAD⁺ rises with age despite declines in selected biosynthetic enzymes
- Mid-adult NAD⁺ labeling declines; NA-derived NR labeling becomes undetectable
- Early Complex I remodeling accompanies altered redox and energetic status
- Loss of NAD⁺ consumers or salvage enzymes reduces reproduction and lifespan

Graphical Abstract

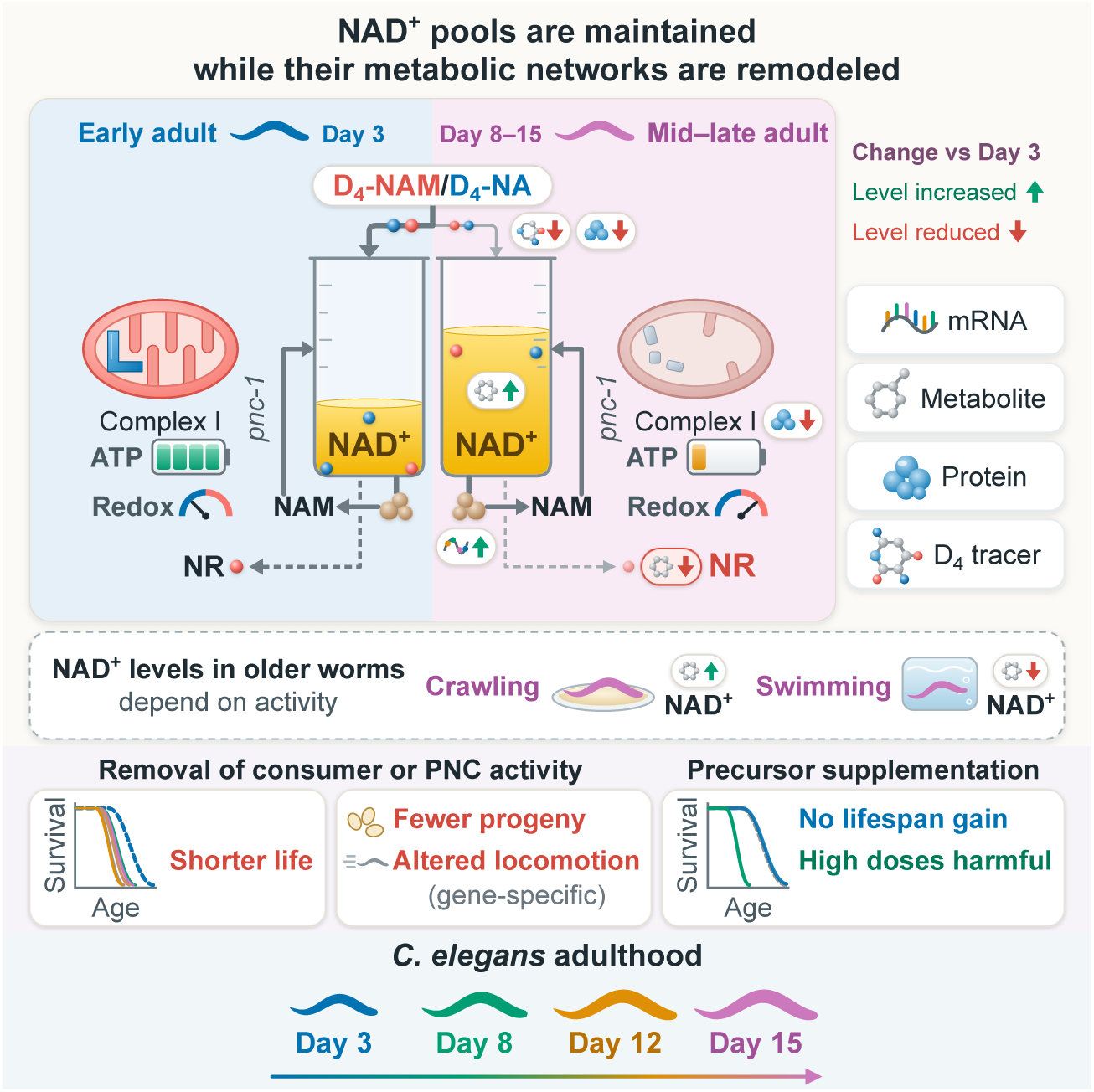

## Introduction

Nicotinamide adenine dinucleotide (NAD⁺) connects cellular bioenergetics with regulatory signaling. As a redox coenzyme, it supports glycolysis **[1, 2]**, the tricarboxylic acid cycle **[3, 4]**, oxidative phosphorylation **[5]**, and other central metabolic pathways, including fatty acid β-oxidation and amino acid metabolism **[6–8]**. NAD⁺ also serves as a substrate for signaling enzymes, including poly(ADP-ribose) polymerases (PARPs) **[9]**, NADases such as CD38 and CD157 **[10–14]**, sirtuins **[15]**, and sterile alpha and Toll/interleukin-1 receptor motif-containing protein 1 (SARM1) **[16]**. Coordination between the metabolic and signaling roles of NAD⁺ is essential for maintaining NAD⁺ homeostasis. Age-associated changes in NAD⁺ abundance have been reported in rodents **[17]**, nematodes **[18]**, and humans **[19]** and have been linked to cellular senescence **[20–23]**, glucose intolerance **[19, 24–27]**, mitochondrial dysfunction **[17, 18, 28–30]**, and neurodegenerative disease **[31–34]**. Understanding how NAD⁺ metabolism and homeostasis changes with age is thus essential to defining its contribution to physiological decline.

Age-related reductions in NAD⁺ have commonly been attributed to diminished biosynthesis and increased consumption in response to chronic stress, including PARP activation during DNA damage **[6, 30, 35–38]** and NADase activity associated with inflammation **[12, 13, 17, 21, 39, 40]**. These observations have motivated efforts to restore NAD⁺ abundance through precursor supplementation **[41–44]**. However, steady-state NAD⁺ abundance reflects the balance between synthesis and consumption and cannot independently resolve the rates of either process or the recycling of precursors. A maintained NAD⁺ pool may therefore conceal substantial changes in the pathways that sustain it, just as a smaller pool does not necessarily indicate reduced synthesis. Indeed, work in aged mice demonstrated that NAD⁺ synthesis flux is maintained despite lower tissue concentrations, with faster fractional turnover in affected tissues **[25]**. Because biosynthetic pathways and NAD⁺-consuming enzymes draw on interconnected metabolite pools, changes in one branch can influence substrate availability and activity elsewhere in the network **[20, 24, 37, 45, 46]**. Yet how these branches are remodeled across the lifespan, and how that remodeling relates to redox balance, energetic status, and organismal function, remains unresolved.

The nematode *Caenorhabditis elegans* provides a tractable model for addressing this gap. Its short lifespan **[47, 48]** enables sampling across successive stages of adulthood, while its sequenced genome **[49]**, conserved metabolic pathways, and genetic accessibility support functional analysis **[50–53]**. Its NAD⁺ biosynthetic architecture also provides a useful context for examining precursor recycling. Mammals salvage nicotinamide (NAM) through nicotinamide phosphoribosyltransferase (NAMPT) **[54–58]** and synthesize NAD⁺ through the *de novo* kynurenine pathway from tryptophan **[59–61]** and the Preiss–Handler pathway from nicotinic acid (NA) **[62]**. *C. elegans* lacks NAMPT and the major mammalian NADase CD38 **[11, 63–67]** but retains the kynurenine pathway **[59–61]** and recycles NAM through the nicotinamidase PNC-1, which converts NAM to NA for entry into the Preiss–Handler pathway [63, 68]. Conserved NAD⁺ consumers include PARPs encoded by *parp-1* and *parp-2* **[50, 69, 70]**, the SARM1 ortholog TIR-1 **[71–73]**, and sirtuins encoded by *sir-2.1* through *sir-2.4* **[74, 75]**. Together, these features enable investigation of how biosynthesis, precursor recycling, and NAD⁺ consumption contribute to metabolic remodeling and physiological function during aging.

Here, we examined NAD⁺ network remodeling across the *C. elegans* adult lifespan by integrating liquid chromatography-mass spectrometry (LC-MS) metabolomics, D₄-NAM and D₄-NA (deuterium) stable-isotope tracing, gene-expression profiling, and proteomics. This approach linked age-dependent changes in NAD⁺ abundance and precursor incorporation with the expression of network components, redox balance, and energetic status. We complemented these analyses with genetic disruption of NAD⁺-consuming and salvage enzymes and supplementation with NAM, NA, and trigonelline (Trigo) **[76]** to assess consequences for locomotion, reproduction, and lifespan. We found that steady-state NAD⁺ abundance was maintained or increased across adulthood despite reductions in several biosynthetic components and diminished precursor incorporation during mid-adulthood. These changes coincided with altered redox and energetic status and early remodeling of mitochondrial Complex I protein abundance. Genetic disruptions produced distinct physiological defects and shortened lifespan, while supplementation outcomes depended on metabolite identity and concentration. Together, these findings reveal extensive age-dependent remodeling of the NAD⁺ network that is not captured by steady-state NAD⁺ abundance alone.

## Results

### Age-dependent transcriptional remodeling distinguishes NAD⁺ biosynthetic and consuming pathways

To examine NAD⁺ network remodeling across adulthood under basal conditions, we established synchronized *C. elegans* cohorts without reproductive inhibitors, such as 5-fluoro-2′-deoxyuridine (FUdR) **[77]**, or sterilizing mutations. Animals were synchronized by physical microfiltration using NemaSync, followed by daily gravity-based separation to maintain adult cohorts free of progeny. Samples were collected at 62 hours after egg laying (D03) and on Days 8 (D08), 12 (D12), and 15 (D15), spanning distinct stages of adulthood (**Figure 1A**).

**Figure 1.**
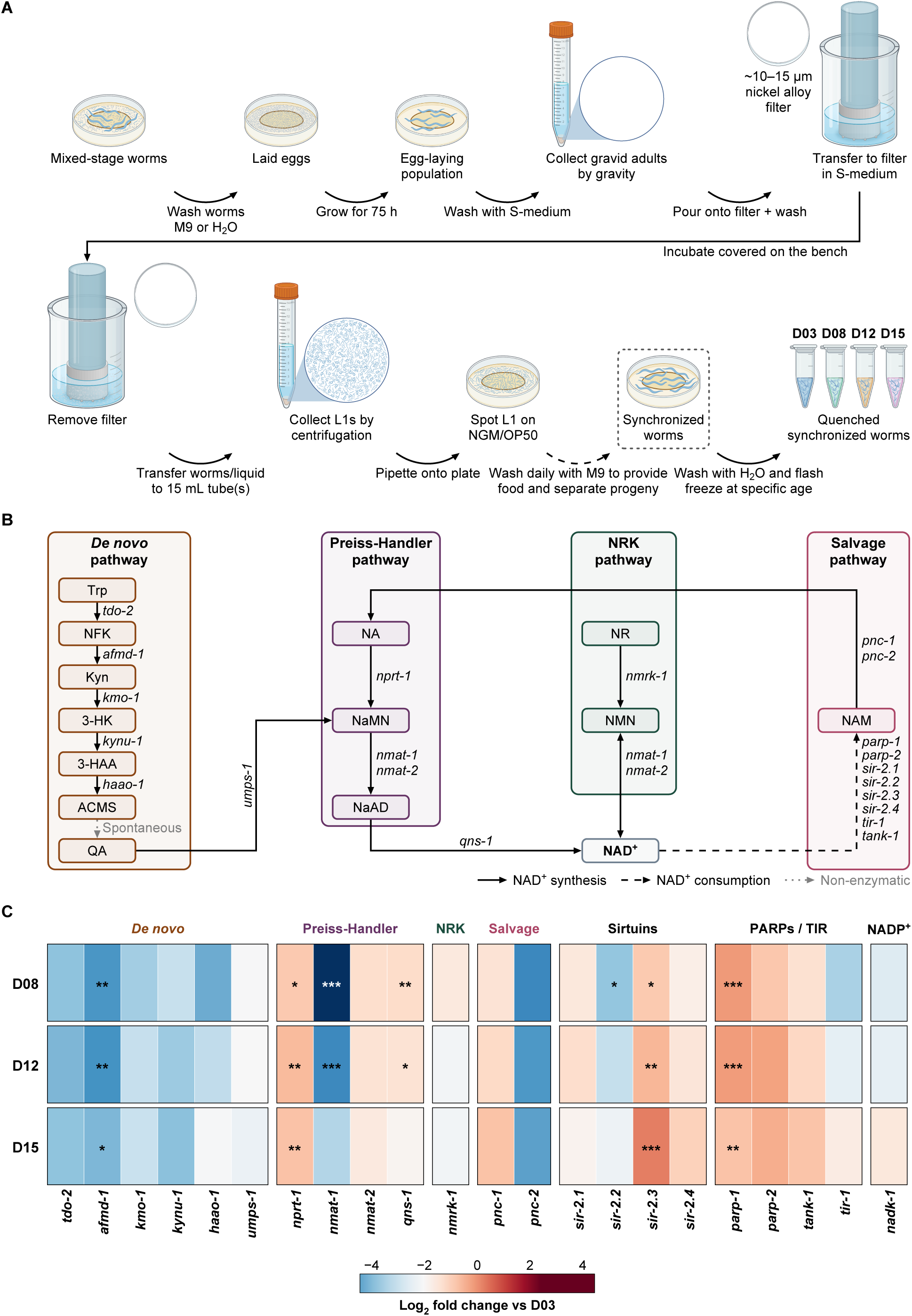
| Age-dependent transcriptional remodeling distinguishes NAD⁺ biosynthetic and consuming pathways. **(A)** Microfiltration and gravity-based separation workflow used to establish synchronized *C. elegans* cohorts and maintain adults free of progeny without reproductive inhibitors or sterilizing mutations. Animals were collected at 62 h after egg laying (D03) and on Days 8 (D08), 12 (D12), and 15 (D15) for molecular profiling. **(B)** Organization of the *C. elegans* NAD⁺ metabolic network, including the *de novo* kynurenine pathway (*tdo-2, afmd-1, kmo-1, kynu-1,* and *haao-1*), the Preiss–Handler pathway (*nprt-1, nmat-1, nmat-2,* and *qns-1*), nicotinamide riboside salvage through *nmrk-1*, and deamidated nicotinamide salvage through *pnc-1* and *pnc-2*, which feeds into the Preiss–Handler pathway. The kynurenine pathway includes spontaneous cyclization of 2-amino-3-carboxymuconate semialdehyde (ACMS) to quinolinate. NAD⁺ consumers that release nicotinamide include the poly(ADP-ribose) polymerases *parp-1* and *parp-2*, tankyrase *tank-1*, sirtuins *sir-2.1* through *sir-2.4*, and the SARM1-like NADase *tir-1*. Solid arrows indicate biosynthetic reactions, dashed arrows indicate NAD⁺ consumption, and dotted gray arrows indicate nonenzymatic reactions. Canonical NAMPT and CD38 orthologs are absent from the *C. elegans* genome. **(C)** Relative expression of 22 NAD⁺-related genes at D08, D12, and D15, measured by quantitative reverse-transcription PCR. Heatmap colors represent mean log₂ fold changes relative to D03; genes are ordered by their D15 fold change. Expression was normalized to the geometric mean of three reference genes: *cdc-42, cyc-1,* and *pmp-3*. Data represent six independent biological replicates per age. For each gene, age groups were compared using one-way ANOVA followed by Dunnett’s multiple-comparison test against D03. *p < 0.05, **p < 0.01, and ***p < 0.001; no asterisk indicates p ≥ 0.05.

The *C. elegans* NAD⁺ network connects *de novo* synthesis through the kynurenine pathway with precursor salvage through the Preiss–Handler, nicotinamide, and nicotinamide riboside (NR) pathways. NR salvage involves NR kinases conserved from yeast to mammals **[55–57]**, whereas nicotinamide salvage feeds the Preiss–Handler pathway through conversion of nicotinamide to nicotinic acid. NAD⁺-consuming enzymes, including sirtuins, poly(ADP-ribose) polymerases, tankyrase, and the SARM1-like NADase TIR-1, release nicotinamide for potential recycling (**Figure 1B**). To assess transcriptional changes across the NAD⁺ network, we quantified the expression of all 22 genes shown in the network schematic (**Figure 1B**) using quantitative reverse-transcription PCR (**Figure 1C**). Pathway-grouped profiles and replicate-level fold changes are shown in **Figure S1A** and **S1B**, respectively (*n* = 6 biological replicates per age).

Selected biosynthetic transcripts declined with age (**Figure 1C and S1**). Expression of the *de novo* pathway gene *afmd-1*decreased significantly at D08 and remained lower at subsequent ages. *tdo-2* expression showed log₂ fold changes of approximately −1.7 to −2.2 relative to D03, although these differences were not significant. The downstream genes *kmo-1*, *kynu-1*, and *haao-1* also showed nonsignificant numerical decreases, whereas *umps-1* expression did not change significantly. The largest reduction occurred in *nmat-1*, which encodes a nicotinamide/nicotinic acid mononucleotide adenylyltransferase. Its expression decreased by more than 4 log₂ units at D08 and partially recovered thereafter, remaining significantly below D03 at D12 but not at D15 (**Figure S1B**).

Other biosynthetic components followed different trajectories. Within the Preiss–Handler pathway, *nprt-1* expression increased by approximately 1.2–1.4 log₂ units and was significantly elevated at all three later ages. *qns-1* expression increased modestly but significantly at D08 and D12 (**Figure S1A and S1B**). Neither the nicotinamide salvage genes *pnc-1*and *pnc-2* nor the NR kinase gene *nmrk-1* changed significantly, although *pnc-2* showed numerical decreases exceeding 2 log₂ units at each later age. Expression of the NAD⁺ kinase gene *nadk-1* also did not change significantly (**Figure S1B**).

Among NAD⁺ consumers, *parp-1* and *sir-2.3* were significantly elevated at all three later ages (**Figure 1C and S1B**). *parp-1* expression increased by more than 1.5 log₂ units at D08 and D12 and remained elevated at D15, whereas *sir-2.3* increased progressively to its highest level at D15. Other consumer genes showed more variable responses: numerical increases in *parp-2*, *tank-1*, and *sir-2.4* did not reach significance, and *tir-1* did not differ significantly from D03. The only significant reduction among consumer transcripts was a transient decrease in *sir-2.2* at D08 (**Figure S1A and S1B**).

Together, these findings reveal selective transcriptional remodeling rather than a uniform decline in NAD⁺ biosynthesis or increase in consumption. Reduced *afmd-1* and *nmat-1* expression coexisted with increased expression of Preiss–Handler components and selected NAD⁺ consumers. Because transcript abundance alone does not establish protein abundance or pathway activity, we next examined age-dependent changes in the NAD⁺-associated proteome.

### The NAD⁺-associated proteome declines broadly with distinct age-dependent trajectories

To determine whether transcriptional remodeling of the NAD⁺ network was accompanied by changes in protein abundance, we performed shotgun proteomic profiling at D03, D08, D12, and D15. We quantified 124 proteins involved in NAD⁺ metabolism or NAD⁺-dependent biochemical processes and grouped them into 12 functional categories (**Figure S2D**).

Reductions in protein abundance became more widespread with age across functional categories. Relative to D03, 23 proteins were significantly reduced at D08, 41 at D12, and 72 at D15, whereas only one, two, and two proteins, respectively, were significantly increased (FDR-adjusted *q* < 0.05 and |log₂ fold change| > 1; **Figures 2A**, **S2A**, and **S2B**). Although median log₂ fold changes remained near zero in most categories at D08, all categories had negative medians by D15 (**Figure S2D**). The largest median reductions occurred in fatty acid synthesis, NADPH production, and amino acid metabolism. The few significantly increased proteins belonged to categories encompassing glycolysis and NADH shuttles, amino acid metabolism, and sugar and other metabolic processes (**Figure S2D**). By D15, median protein abundance was lower than at D03 in all 12 NAD⁺-associated functional categories, revealing a broad decline across the measured proteome.

Principal component analysis resolved age-associated differences in protein profiles (**Figure 2B**). The first principal component explained 54.9% of the total variance and broadly ordered samples by age, with D03 samples forming a distinct cluster followed by D08, D12, and D15. Clustering identified groups of proteins that declined early in adulthood (cluster 1) or at later ages (cluster 2) (**Figure 2C**), revealing distinct temporal trajectories within the broader proteomic decline.

**Figure 2.**
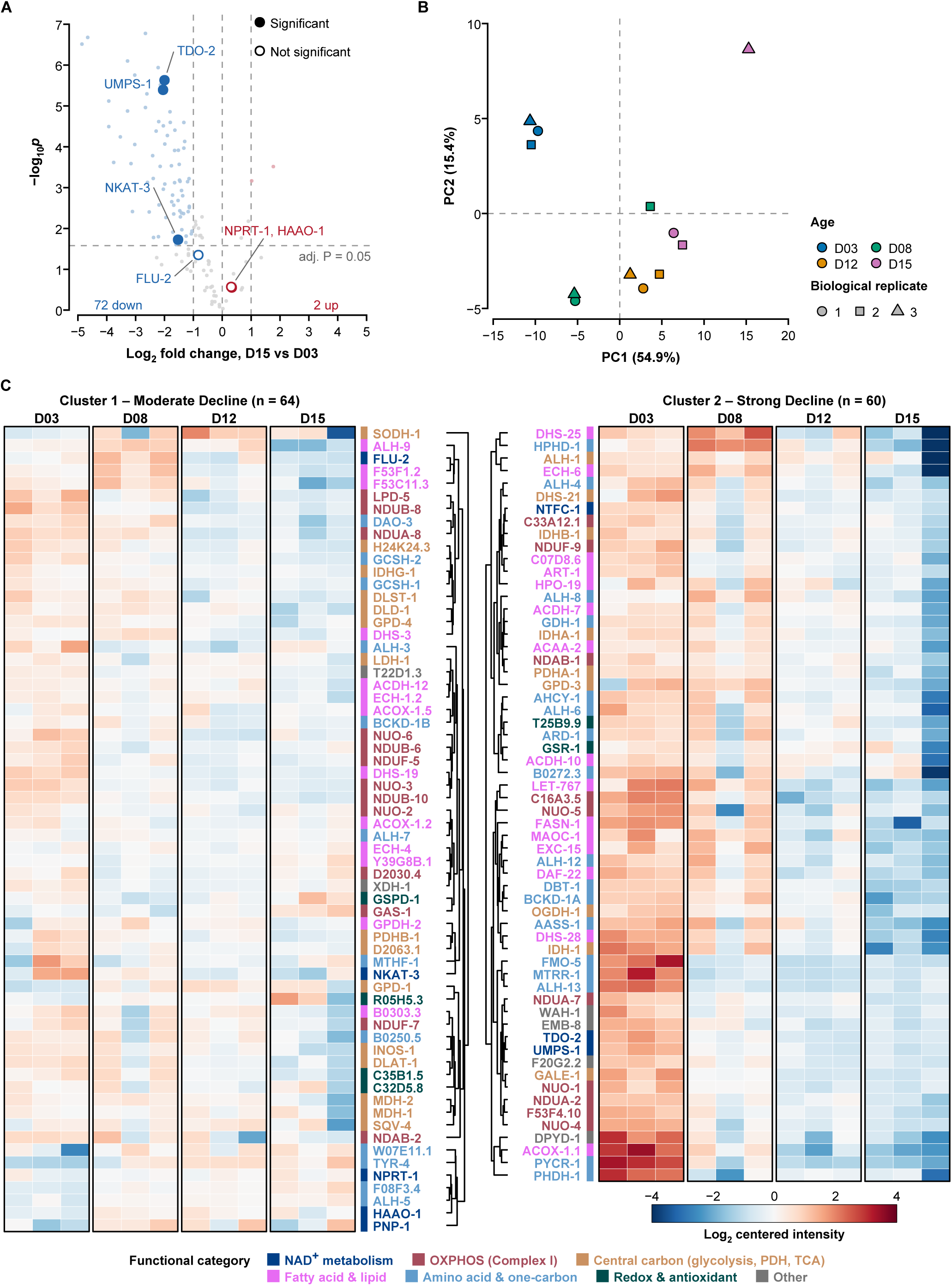
| The NAD⁺-associated proteome declines broadly with distinct age-dependent trajectories. **(A)** Volcano plot of protein-abundance changes at D15 relative to D03 for 124 NAD⁺-related and NAD⁺-dependent proteins. Vertical dashed lines indicate the fold-change threshold (|log₂ fold change| > 1), and the horizontal dashed line indicates the Benjamini–Hochberg FDR-adjusted significance threshold (*q* < 0.05). Filled circles indicate proteins meeting both criteria; open circles indicate proteins that did not meet the combined criteria. Seventy-two proteins met the criteria for decreased abundance. Only two proteins exhibited positive fold changes, neither of which met the combined criteria. **(B)** Principal component analysis of the 124 proteins across D03, D08, D12, and D15. Protein intensities were log₂-transformed and autoscaled before analysis. PC1 and PC2 explain 54.9% and 15.4% of the total variance, respectively. PC1 broadly orders samples by age, with D03 and D15 at opposite ends of the axis. Symbol color indicates age, and symbol shape identifies the biological replicate. **(C)** Heatmap of row-centered log₂ protein intensities across 12 samples, representing three independent biological replicates per age. *k*-means clustering (*k* = 2) identified 64 proteins that declined beginning in early adulthood and 60 proteins that remained comparatively stable until D15. Within each cluster, proteins were ordered by hierarchical clustering using Euclidean distance and complete linkage. Colors indicate abundance relative to each protein’s mean log₂ intensity, ranging from −4 (blue) to +4 (warm colors); white indicates the row mean. Data are presented as mean ± SD from three independent biological replicates per age. For differential-abundance analyses in (A) statistical significance was assessed using moderated two-sided *t* tests with Benjamini–Hochberg FDR correction across all 1,810 quantified proteins. Proteins were classified as altered when *q*< 0.05 and |log₂ fold change| > 1.

Selected enzymes associated with NAD⁺ biosynthesis also showed early or progressive reductions (**Figures 2A** and **S2A–S2C**). TDO-2 and UMPS-1 abundance decreased significantly at D08 and remained lower through D15, whereas NKAT-3 declined more gradually and reached significance only at D15. FLU-2 (KYNU-1), HAAO-1, and NPRT-1 did not meet the criteria for significant change at any sampled age (**Figure S2C**). FLU-2 increased numerically at D08 before declining at D12 and D15, HAAO-1 showed a modest numerical increase, and NPRT-1 remained near D03 levels.

Protein trajectories did not consistently parallel transcript changes. Despite significantly increased *nprt-1* expression, NPRT-1 abundance remained comparatively stable. Likewise, HAAO-1 abundance did not mirror the numerical decrease in *haao-1* transcript at D08 (**Figures S1B** and **S2C**). These observations highlight differences between transcript and protein profiles without establishing the regulatory mechanisms underlying them.

Together, these findings reveal increasingly widespread reductions in the NAD⁺-associated proteome, with distinct trajectories across functional groups and individual biosynthetic components. Because protein abundance does not directly establish pathway activity, these changes leave unresolved whether the NAD⁺ pool declines alongside the proteins that support its metabolism. We therefore next examined steady-state NAD⁺ abundance, its precursors, and related redox metabolites across adulthood.

### Steady-state NAD⁺ abundance increases amid stage-specific remodeling of the NAD⁺ metabolome

To determine whether transcriptional and proteomic remodeling was accompanied by changes in metabolite abundance, we profiled the steady-state NAD⁺ metabolome at D03, D08, D12, and D15. Using liquid chromatography–mass spectrometry (LC-MS), we measured NAD⁺, related cofactors, intermediates of both *de novo* and salvage biosynthesis, kynurenine-pathway side-branch metabolites, and products associated with NAD⁺ consumption and clearance (**Figure S3B**).

Partial least-squares discriminant analysis revealed age-associated differences in the targeted metabolomic profiles (**Figure 3A**). The first two components explained 27.9% and 21.1% of the variance, respectively. D03 samples separated from later ages primarily along the first component, whereas D08 samples separated mainly along the second. D12 and D15 profiles largely overlapped on these two components. Variable importance in projection (VIP) scores summarize how strongly each metabolite contributes to the model’s separation of age groups; scores above 1 indicate above-average contributions. Ten metabolites exceeded this threshold, with the highest scores for nicotinamide mononucleotide (NMN), 3-hydroxykynurenine (3-HK), 3-hydroxyanthranilic acid (3-HAA), tryptophan (Trp), and NAD⁺ (**Figure S3A**). These metabolites span the *de novo* kynurenine pathway, the NR salvage pathway, and the NAD⁺ pool itself, indicating that differences across the biosynthetic network contributed to the separation. NADH and NADP⁺ had the lowest VIP scores, indicating that they contributed least to distinguishing age groups in this model.

**Figure 3.**
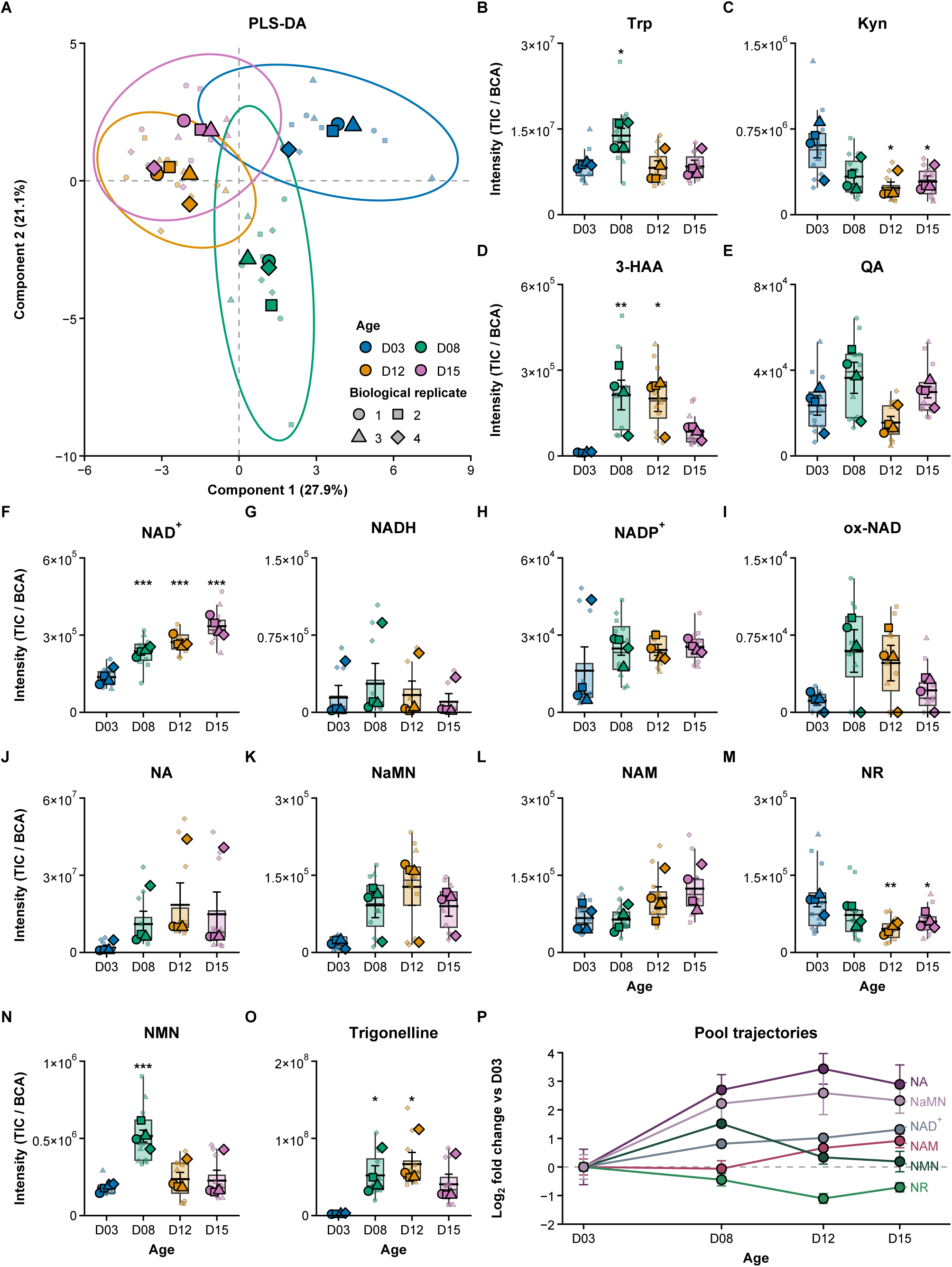
| Steady-state NAD⁺ abundance increases amid stage-specific remodeling of the NAD⁺ metabolome. **(A)** Partial least-squares discriminant analysis (PLS-DA) score plot of 24 targeted NAD⁺-related metabolites measured at 62 h after egg laying (D03) and on Days 8 (D08), 12 (D12), and 15 (D15). The model used 16 independent biological samples, with technical injections averaged within each sample. Components 1 and 2 explain 27.9% and 21.1% of the modeled variance, respectively. Model performance was evaluated by cross-validation and permutation testing (R²Y = 0.62, Q² = 0.41, permutation *p* ≤ 0.005). Ellipses represent 95% bivariate normal distributions and illustrate within-group dispersion. **(B–E)** Normalized abundance of *de novo* kynurenine-pathway metabolites: tryptophan (B), kynurenine (C), 3-hydroxyanthranilic acid (3-HAA; D), and quinolinate (QA; E). Tryptophan increased transiently at D08, whereas kynurenine was significantly lower at D12 and D15 than at D03. 3-HAA increased at D08 and D12 before declining at D15. Quinolinate reached its highest abundance at D08 and decreased at D12 but these differences were not significant. **(F–I)** Normalized abundance of NAD⁺ (F), NADH (G), NADP⁺ (H), and ox-NAD (I). NAD⁺ abundance increased across adulthood (overall age effect, *q* = 2.7 × 10⁻⁵), whereas NADH (*q* = 0.8325) and NADP⁺ (*q* = 0.5743) did not differ significantly across ages. Ox-NAD showed a numerical increase at D08 and D12 followed by a decline at D15, but the overall age effect was not significant. **(J–O)** Normalized abundance of nicotinic acid (NA; J), nicotinic acid mononucleotide (NaMN; K), nicotinamide (NAM; L), nicotinamide riboside (NR; M), nicotinamide mononucleotide (NMN; N), and trigonelline (O). NA (*q* = 0.4192) and NAM (*q* = 0.1034) did not differ significantly across ages. NaMN increased through D12 before declining at D15. NMN increased at D08 and subsequently returned toward D03 abundance (overall age effect, *q* = 0.0053). NR declined significantly with age, reaching its lowest abundance at D12 and remaining lower at D15. Trigonelline increased at D08 and D12 before declining at D15. **(P)** Group-mean trajectories of NAD⁺, NAM, NMN, NA, NaMN, and NR, expressed as log₂ fold changes relative to D03. Divergent trajectories highlight D08–D12 as an interval of pronounced metabolite remodeling, including reduced NR abundance. For (B–O), metabolite signals were normalized to total ion count and BCA-measured protein abundance. Boxplots show medians and interquartile ranges (IQRs), with whiskers extending to the most extreme observations within 1.5 × IQR of the box boundaries. Large, outlined symbols represent the mean of technical injections for each biological replicate; smaller translucent symbols represent individual injections. Statistical analyses used four independent biological replicates per age. Overall age effects were assessed by one-way ANOVA, with Benjamini–Hochberg FDR correction across the 14 metabolites in (B–O). Pairwise comparisons were assessed using Tukey’s honestly significant difference test. Only comparisons with the corresponding D03 group are displayed. Asterisks indicate significant differences relative to D03: *p < 0.05, **p < 0.01, and ***p < 0.001; unmarked comparisons were not significant.

Metabolites within the *de novo* kynurenine pathway followed distinct temporal patterns (**Figures 3B–3E** and **S3B–S3D**). Trp increased transiently at D08 before returning toward D03 levels at D12 and D15 (**Figure 3B**). N-formylkynurenine (NFK) declined numerically after D03, but the differences were not significant (**Figure S3C**). Kynurenine (Kyn) decreased and was significantly lower at D12 and D15 than at D03 (**Figure 3C**). In contrast, 3-HK increased significantly at D08 before returning toward D03 levels (**Figure S3D**), while 3-HAA increased significantly at D08 and D12 before declining at D15 (**Figure 3D**). Quinolinate was numerically highest at D08 and lower at D12, although these differences were not significant (**Figure 3E**). Thus, steady-state levels of *de novo* pathway intermediates did not decline uniformly alongside the reductions in selected biosynthetic transcripts and proteins.

Kynurenine-pathway side-branch metabolites also exhibited heterogeneous, nonsignificant changes (**Figures S3B** and **S3E–S3I**). Kynurenic acid and picolinic acid increased numerically across adulthood, reaching their highest abundances at D12–D15 (**Figure S3E** and **S3I**). 4-(2-Amino-3-hydroxyphenyl)-2,4-dioxobutanoate (AHPDB) and 2-aminomuconic 6-semialdehyde (2-AMS) were numerically lowest at D08 before increasing at later ages, whereas 2-aminomuconate peaked at D08 (**Figure S3F–S3H**). These profiles suggest possible changes in the distribution of metabolites among kynurenine-pathway branches, but they do not establish increased diversion of flux away from quinolinate formation.

Despite reductions in several biosynthetic components, NAD⁺ abundance increased progressively across adulthood and was highest at D15 (**Figure 3F**). NADH and NADP⁺ did not differ significantly across the sampled ages (**Figure 3G** and **3H**). Overoxidized NAD (ox-NAD) increased numerically at D08 and D12 before declining at D15, but these differences were not significant (**Figure 3I**). The steady-state NAD⁺ pool therefore did not mirror the age-dependent decline in selected biosynthetic transcripts and proteins.

Precursors and salvage-associated metabolites likewise followed distinct trajectories (**Figures 3J–3O** and **S3J**). Nicotinic acid (NA) and nicotinamide (NAM) did not differ significantly across adulthood despite numerical increases at later ages (**Figure 3J** and **3L**). Nicotinic acid mononucleotide (NaMN) and nicotinic acid riboside (NaR) increased numerically through D12 without reaching significance (**Figures 3K** and **S3J**). NMN increased sharply at D08 and subsequently returned toward D03 levels (**Figure 3N**), whereas NR was significantly reduced at D12 and D15 (**Figure 3M**). Trigonelline increased at D08 and D12 before declining at D15 (**Figure 3O**). Cyclic ADP-ribose (cADPR) remained near D03 levels at D08, increased at D12, and was significantly elevated at D15, coinciding with peak NAD⁺ abundance (**Figure S3K**). Comparison of normalized metabolite trajectories highlighted D08–D12 as an interval of pronounced remodeling: NA and NaMN reached their maxima within this interval, NMN returned toward baseline, and NR fell to its lowest abundance (**Figure 3P**).

Together, these results reveal divergent trajectories among the NAD⁺ pool, its precursors, side-branch metabolites, and consumption-associated products. NAD⁺ accumulated across adulthood despite reductions in selected biosynthetic transcripts and proteins, while NR emerged as a precursor with a significant age-associated decline. Because steady-state measurements cannot distinguish changes in production, utilization, recycling, or clearance, they leave unresolved how precursor handling contributes to maintenance of the NAD⁺ pool. We therefore next used *in vivo* D₄-nicotinamide and D₄-nicotinic acid tracing to examine precursor incorporation into NAD⁺ and related metabolites.

### *In vivo* isotope tracing reveals age-dependent changes in NAD⁺ maintenance and precursor-specific loss of NR labeling

To complement our steady-state measurements and obtain a more complete picture of how NAD⁺ metabolism changes with age, we used *in vivo* stable-isotope tracing to examine how worms incorporate and process NAD⁺ precursors. Synchronized animals at D03, D08, D12, and D15 were incubated with either D₄-nicotinamide (D₄-NAM) or D₄-nicotinic acid (D₄-NA) for four hours in liquid culture with continuous rocking. Worms and extracellular media were collected separately for LC-MS analysis (**Figure 4A**). We used isotopologue distributions to assess precursor incorporation into NAD⁺ and related metabolites, identify labeling patterns consistent with NAD(H) redox cycling, and track the appearance of labeled metabolites in the extracellular medium (**Figures 4B** and **S4**).

**Figure 4.**
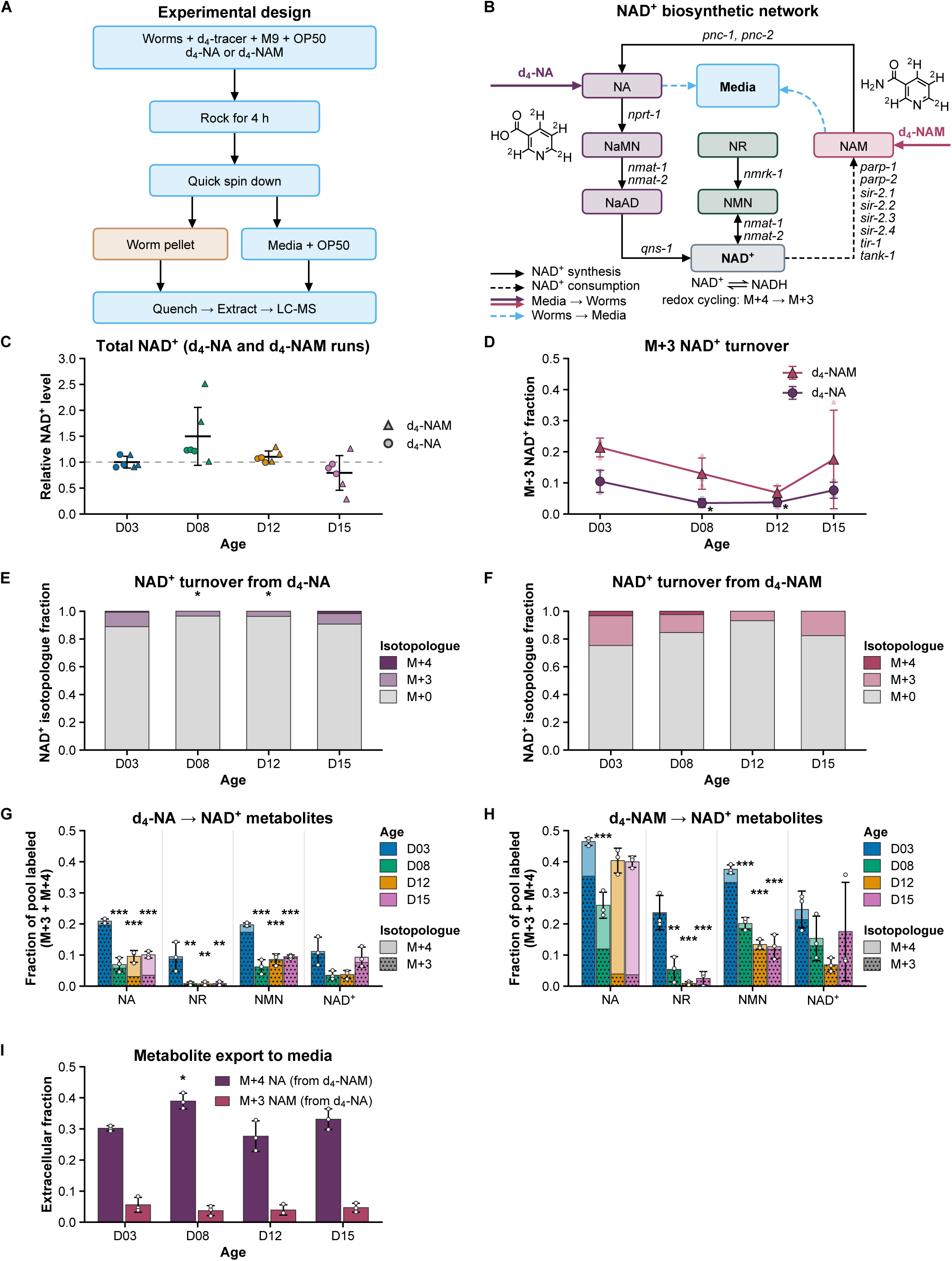
| *In vivo* isotope tracing reveals age-dependent changes in NAD⁺ maintenance and precursor-specific reductions in NR labeling. (A) Stable-isotope pulse-labeling workflow. Synchronized *C. elegans* cohorts at 62 h after egg laying (D03) and on Days 8 (D08), 12 (D12), and 15 (D15) were incubated with D₄-nicotinamide (D₄-NAM) or D₄-nicotinic acid (D₄-NA) for 4 h in liquid culture with continuous rocking. Worms were separated from the extracellular medium by centrifugation, quenched, and extracted for LC-MS analysis. Worm pellets and media were analyzed separately. **(B)** Schematic of tracer incorporation into the NAD⁺ network. D₄-NA enters the Preiss–Handler pathway, whereas D₄-NAM is first deamidated to labeled NA by PNC-1 and PNC-2 before entering the shared downstream pathway. Incorporation of either D₄ precursor generates M+4 NAD⁺; loss of a labeled deuterium during NAD⁺/NADH redox cycling can generate M+3 NAD⁺. **(C)** Relative total NAD⁺ abundance during the tracing experiments. Values from both tracer conditions were pooled by age and normalized to the pooled D03 mean (dashed line, 1.0). NAD⁺ abundance increased numerically at D08 and declined at D12 and D15, but the overall age effect was not significant. This pattern contrasts with the progressive NAD⁺ accumulation observed under plate-culture conditions (Figure 3F). Horizontal lines indicate means, and whiskers indicate ± SD. Data represent six independent biological replicates per age, comprising three replicates per tracer condition. Symbol shape identifies the tracer condition. **(D)** Fractional enrichment of M+3 NAD⁺ following D₄-NAM or D₄-NA exposure. NAM-derived enrichment was greater than NA-derived enrichment across the sampled ages. Enrichment from both tracers decreased at D08 and D12 relative to D03 and partially recovered at D15. **(E and F)** NAD⁺ mass-isotopologue distributions following D₄-NA (E) or D₄-NAM (F) exposure. M+0 represents unlabeled NAD⁺, M+4 represents NAD⁺ retaining all four tracer-derived deuteriums, and M+3 represents NAD⁺ retaining three deuteriums, consistent with label loss during redox cycling. Asterisks indicate significant differences in the M+3 fraction relative to D03 within each tracer condition. (**G and H)** Intracellular labeling of NA, NR, NMN, and NAD⁺ following D₄-NA (G) or D₄-NAM (H) exposure. Stacked bars show M+3 and M+4 fractions; total bar height represents their combined labeled fraction. With D₄-NA, NA and NMN labeling decreased significantly at D08, D12, and D15 relative to D03, NR labeling became undetectable at all three later ages, and NAD⁺ labeling decreased at D08 and D12 (G). With D₄-NAM, NA labeling decreased significantly at D08, while NR and NMN labeling decreased significantly at D08, D12, and D15. Low residual NR labeling remained at D08 and D15, and NAD⁺ labeling showed a downward trend (H). Asterisks indicate significant differences in the combined labeled fraction relative to D03 for each metabolite. **(I)** Fractional enrichment of labeled metabolites in the extracellular medium. Following D₄-NAM exposure, M+4 NA represented approximately 30–40% of the extracellular NA pool. Following D₄-NA exposure, M+3 NAM represented less than 10% of the extracellular NAM pool. These values describe enrichment of the respective metabolite pools, rather than the fraction of administered tracer released into the medium. For (D–I), data are presented as mean ± SD from three independent biological replicates per age and tracer condition. Age-associated differences were evaluated separately for each tracer and metabolite using one-way ANOVA followed by Tukey’s honestly significant difference test. Only comparisons with the corresponding D03 group are displayed. *p < 0.05, **p < 0.01, and ***p < 0.001; unmarked comparisons were not significant.

Both tracers carry deuterium at C2, C4, C5, and C6 of the pyridine ring, yielding M+4 NAD⁺ upon incorporation. Loss of the C4 deuterium during NAD(H) redox cycling generates M+3 NAD⁺ **[78]**. Subsequent NAD⁺ consumption releases M+3 NAM, which can be converted to M+3 NA through nicotinamidase activity and re-enter biosynthetic pathways to regenerate labeled NAD⁺(M+3), linking redox turnover to consumer-mediated precursor recycling (**Figure S4**).

Total NAD⁺ abundance followed a different age-dependent trajectory under tracing conditions than on standard NGM plates. Whereas NAD⁺ increased progressively across adulthood on plates (**Figure 3F**), animals incubated in liquid with continuous rocking showed an increase at D08 followed by lower abundance at D12 and D15 (**Figure 4C**). This numerical pattern occurred with both tracers, although the age-associated differences were not significant. Older animals therefore did not show the progressive NAD⁺ accumulation observed under plate conditions. Sustained swimming may impose greater energetic demand than plate culture, where animals can move intermittently and potentially conserve energy. However, the contributions of movement, energy expenditure, and other differences between the culture conditions cannot be distinguished by these experiments.

Under matched labeling conditions, D₄-NAM yielded greater M+3 NAD⁺ enrichment than D₄-NA at all four ages (**Figure 4D**). NAM enters the Preiss–Handler pathway after conversion to NA by the nicotinamidases PNC-1 and PNC-2 (**Figure S4**). Thus, despite requiring this additional step, NAM contributed more label to the M+3 NAD⁺ pool than directly supplied NA. One possibility is that worms take up NAM more readily than NA, making more labeled NA available intracellularly after NAM deamidation. However, M+3 NAD⁺ enrichment alone cannot distinguish differences in uptake from differences in intracellular label dilution or subsequent redox cycling.

NAD⁺ labeling also changed with age. With both tracers, the M+3 fraction decreased at D08, remained low at D12, and partially recovered at D15 (**Figure 4D**). The complete isotopologue distributions similarly showed reduced labeled fractions and a larger unlabeled M+0 contribution during mid-adulthood (**Figure 4E** and **4F**). Together, these measurements identify D08–D12 as an interval of reduced labeling of the NAD⁺ pool from exogenous NAM and NA.

Intracellular metabolites showed precursor-specific labeling patterns, with particularly pronounced changes in NR (**Figure 4G** and **4H**). Following D₄-NA exposure, the labeled fractions of NA and NMN were significantly lower at D08, D12, and D15 than at D03. NR labeling became undetectable at all three later ages, while NAD⁺ labeling decreased at D08 and D12 (**Figure 4G**). Following D₄-NAM exposure, intracellular labeled NA was consistent with nicotinamide deamination, but its labeled fraction decreased significantly only at D08. NR and NMN labeling were significantly lower at all three later ages, whereas NAD⁺ labeling showed a downward trend (**Figure 4H**). Unlike the loss of detectable NR labeling with D₄-NA, low residual NR labeling remained at D08 and D15 following D₄-NAM exposure.

These patterns identify NR-associated metabolism as a particularly age-sensitive feature of the *C. elegans* NAD⁺ network. Loss of detectable NA-derived NR labeling and reduced NAM-derived NR labeling accompanied the significant decline in steady-state NR abundance (**Figure 3M**). Because *C. elegans* lacks a NAMPT ortholog, these labeling patterns cannot be interpreted as direct NAM-to-NMN salvage. Label appearing in NMN and NR may instead involve downstream processing and recycling of labeled NAD⁺ or other network intermediates (**Figure S4**). Reduced labeling could therefore reflect changes in labeled NAD⁺ turnover, NR formation or utilization, or upstream precursor availability.

To examine extracellular precursor handling, we measured the labeled fraction of the NA and NAM pools in the incubation medium (**Figure 4I**). The medium contained autoclaved *E. coli* OP50 as a food source to minimize bacterial metabolism of the tracers. Following D₄-NAM exposure, M+4 NA comprised approximately 30–40% of the extracellular NA pool, consistent with deamination of D₄-NAM and release of labeled NA into the medium (**Figure S4**). In the reverse direction, following D₄-NA exposure, M+3 NAM comprised less than 10% of the extracellular NAM pool, consistent with passage of the label through NAD⁺ biosynthesis, redox cycling, and consumption before NAM release (**Figure S4**). Thus, the extracellular NA pool was more enriched with NAM-derived label than the extracellular NAM pool was with NA-derived label.

Together, these experiments reveal age-dependent changes in precursor handling that are not captured by steady-state NAD⁺ abundance alone. Reduced mid-adult NAD⁺ labeling coincided with complete loss of detectable NA-derived NR labeling and marked reductions in NAM-derived NR labeling. In parallel, the progressive NAD⁺ accumulation observed on plates was absent during liquid incubation with continuous rocking, raising the possibility that age-dependent NAD⁺ maintenance varies with physiological conditions and demand. These findings connect network remodeling to altered precursor utilization and prompted us to examine the accompanying changes in redox balance and energetic status.

### Transient mid-adult ox-NAD accumulation accompanies sustained changes in redox balance and energetic status

To determine whether reduced mid-adult precursor incorporation coincided with broader metabolic changes, we examined ox-NAD/NAD⁺, glutathione and NAD-associated redox ratios, and adenylate energetic indices across the four sampled ages. We also assessed metabolite ratios associated with methylation, carnitine-linked acyl handling, and tricarboxylic acid (TCA) cycle metabolism.

The ox-NAD/NAD⁺ ratio followed a nonlinear trajectory (**Figure 5A**). It increased numerically at D08, remained elevated at D12, and declined at D15. The overall age effect was significant, with both D08 and D12 differing significantly from D15. Thus, ox-NAD abundance relative to NAD⁺ was highest during mid-adulthood rather than accumulating progressively with age.

**Figure 5.**
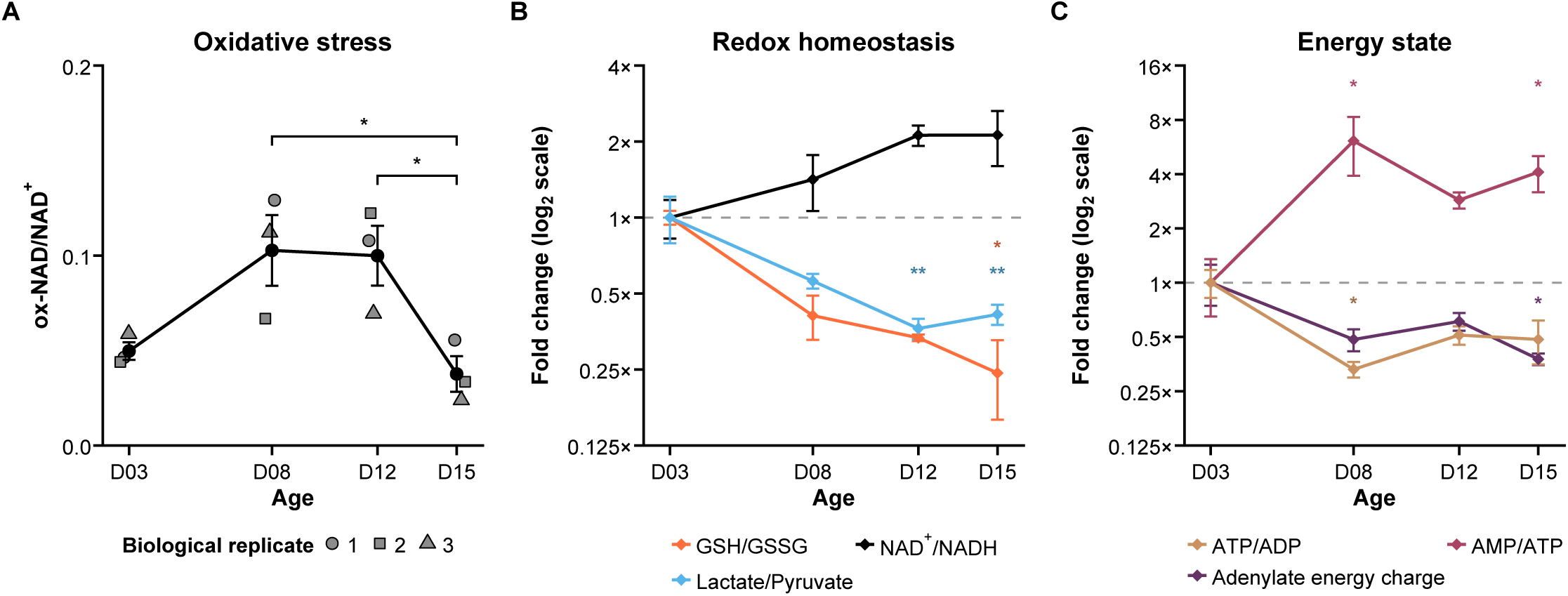
| Transient mid-adult ox-NAD accumulation accompanies sustained changes in redox balance and energetic status. **(A)** Ratio of ox-NAD to NAD⁺ LC-MS peak areas at D03, D08, D12, and D15. The ratio increased numerically at D08 and remained elevated at D12 before declining at D15. The overall age effect was significant (FDR-adjusted *q* = 0.019). Black symbols and connecting lines show mean ± SEM; gray diamonds indicate individual biological replicates. Brackets indicate significant Tukey’s honestly significant difference (HSD) comparisons between D08 and D15 and between D12 and D15. **(B)** Apparent redox indices normalized to the D03 mean and displayed on a log₂ scale. GSH/GSSG (*q* = 0.022) and lactate/pyruvate (*q* = 0.0074) decreased across adulthood, consistent with changes toward a more oxidized metabolic state. NAD⁺/NADH increased numerically, but the overall age effect was not significant (*q* = 0.18). The dashed line indicates the D03 baseline. **(C)** Apparent adenylate energetic indices normalized to the D03 mean and displayed on a log₂ scale. The energy charge index, calculated from peak areas as (ATP + 0.5 × ADP)/(ATP + ADP + AMP), decreased across adulthood (*q* = 0.043). ATP/ADP also decreased (*q* = 0.022), whereas AMP/ATP increased (*q* = 0.022). All three indices were derived from the same full-scan adenylate measurements. The dashed line indicates the D03 baseline. Data are presented as mean ± SEM. Panel A includes four independent biological replicates per age. Panels B and C include three independent biological replicates per age, except NAD⁺/NADH at D12, which includes two. All indices are semi-quantitative LC-MS peak-area ratios and describe relative changes rather than absolute molar ratios; the peak-area-derived energy charge index is not a calibrated molar adenylate energy charge. Measurements represent whole-animal extracts and do not resolve compartment-specific redox states. Overall age effects were assessed by one-way ANOVA of log₂-transformed ratios, followed by Benjamini–Hochberg FDR correction. Pairwise comparisons were assessed using Tukey’s HSD test. Brackets in (A) indicate the displayed pairwise comparisons; color-matched asterisks in (B) and (C) indicate significant differences from D03 for the corresponding index. *p < 0.05 and **p < 0.01.

Other ratios revealed redox changes that persisted into late adulthood (**Figure 5B**). GSH/GSSG decreased progressively, falling by more than 60% by D12 and remaining low at D15, when the reduction reached significance relative to D03. This pattern was consistent with a more oxidized glutathione pool. Lactate/pyruvate likewise declined to approximately 40% of its D03 value and was significantly reduced at D12 and D15. Although this ratio provides an indirect readout of cytosolic NADH/NAD⁺ balance, whole-animal measurements cannot resolve compartment-specific redox states. The measured NAD⁺/NADH ratio increased approximately 2.2-fold, but the difference was not significant. Together, these patterns support a shift toward a more oxidized metabolic state during mid-to-late adulthood.

Adenylate ratios showed an earlier and sustained change (**Figure 5C**). ATP/ADP declined sharply at D08 and remained below D03 levels through D15, whereas AMP/ATP increased by more than fourfold at D08 and remained elevated thereafter. The apparent adenylate energy charge index also decreased significantly beginning at D08 and did not fully recover. Quality-control ratios used to assess ATP breakdown and adenylate kinase equilibrium did not differ significantly across ages (**Figure S5G** and **S5H**), providing no evidence of age-dependent differences in these indicators of sample degradation. These peak-area-derived indices were consistent with lower energetic status despite maintained or increased steady-state NAD⁺ abundance.

Methylation-associated ratios exhibited pronounced mid-adult changes. S-adenosylmethionine/S-adenosylhomocysteine (SAM/SAH) increased sharply at D08 and peaked at D12 before declining at D15, when it remained significantly above D03 (**Figure S5A**). This trajectory reflected both increased SAM and decreased SAH abundance (**Figure S5I**). N¹-methylnicotinamide/nicotinamide (MeNAM/NAM) increased significantly at D08 and D12 before returning toward D03 levels at D15 (**Figure S5E**). Its mid-adult elevation coincided with the peaks in ox-NAD/NAD⁺ and trigonelline abundance (**Figures 5A** and **3O**).

Carnitine– and TCA-associated ratios followed more sustained trajectories. Acetylcarnitine/free carnitine (C2/C0) decreased significantly at D08 and remained low through D15, primarily reflecting free carnitine accumulation rather than acetylcarnitine loss (**Figure S5B** and **S5J**). Succinate/fumarate also decreased significantly beginning at D08, accompanied by a marked increase in fumarate (**Figure S5D** and **S5L**). In contrast, α-ketoglutarate/succinate increased significantly at D12 and D15 alongside α-ketoglutarate accumulation (**Figure S5C** and **S5K**). Glutamine/glutamate did not change significantly (**Figure S5F**). These results identify changes in the relative abundance of metabolites across several pathways without independently establishing changes in pathway flux.

Together, these findings identify D08 as the earliest sampled age showing a pronounced energetic transition accompanied by broader metabolic remodeling. The mid-adult elevations in ox-NAD/NAD⁺ and MeNAM/NAM subsided by D15, whereas altered glutathione redox balance, adenylate energetic indices, C2/C0, and succinate/fumarate persisted. These distinct trajectories show that NAD⁺ accumulation does not coincide with uniform preservation of redox balance or energetic status. We next integrated the metabolomic and proteomic datasets to identify NAD⁺-associated and mitochondrial processes temporally linked to this transition.

### Multi-omic integration reveals divergent NAD⁺ pathway trajectories and early mitochondrial Complex I remodeling

To examine how NAD⁺ network remodeling relates to age-associated changes in redox balance and energetic status, we integrated the metabolomic and proteomic datasets by pathway and functional group. These analyses revealed divergent metabolite trajectories alongside early changes in mitochondrial Complex I proteins.

Metabolites associated with the *de novo* and Preiss–Handler pathways followed opposing trajectories across adulthood (**Figure 6A**). The *de novo* group included Trp, NFK, Kyn, and 3-HK, and the Preiss–Handler-associated group included NA, NaMN, NaR, and NAD⁺ (**Figure S6C**). By D12, the mean log₂ fold change relative to D03 was approximately −1.7 for the *de novo* group and +2.0 for the Preiss–Handler-associated group. NFK and Kyn showed the largest decreases in the *de novo* group, whereas Trp and 3-HK changed less. All four metabolites in the Preiss–Handler-associated group increased by D12, with NA showing the largest increase and NAD⁺ the smallest (**Figure S6C**). The two profiles remained separated at D15, showing that NAD⁺ abundance was maintained or increased even as several *de novo* intermediates declined. These abundance measurements do not establish whether the rate of NAD⁺ synthesis changed through either pathway.

**Figure 6.**
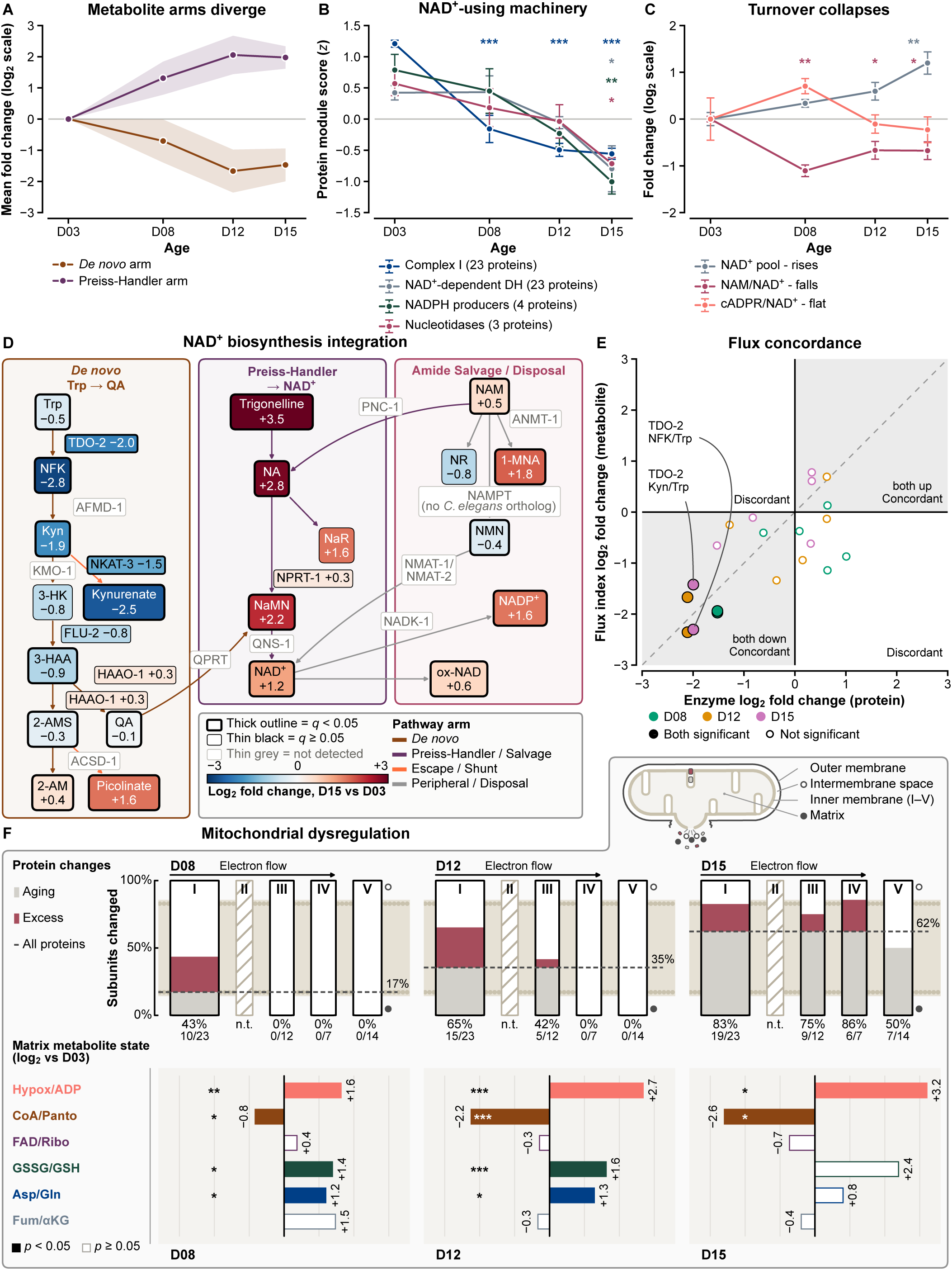
| Multi-omic integration reveals divergent NAD⁺ pathway trajectories and early mitochondrial Complex I remodeling. **(A)** Mean log₂ fold changes relative to D03 for selected metabolites assigned to the *de novo* kynurenine branch or the Preiss–Handler and salvage branch (four metabolites per group). The groups followed opposing trajectories, with approximately 3.5 log₂ units of separation at D12. Shading indicates mean ± SEM across metabolites within each group, rather than variation among biological replicates. These pathway summaries are descriptive and were not used for inferential testing. **(B)** Protein-module z scores at D03, D08, D12, and D15. The Complex I module comprised 23 detected proteins associated with NADH oxidation and declined beginning at D08 (overall age effect, nominal *p* < 0.001). Modules comprising NAD-dependent dehydrogenases (23 proteins; *p* = 0.020), NADPH-producing enzymes (four proteins; *p* = 0.006), and nucleotidases (three proteins; *p* = 0.024) remained near or above their D03 values at D08 before declining later. Reported *p* values describe overall age effects, not individual pairwise comparisons. **(C)** Relative NAD⁺ abundance and consumption-associated metabolite ratios, expressed as log₂ fold changes relative to D03. NAD⁺ abundance increased across adulthood (overall age effect, *p* = 0.007), whereas NAM/NAD⁺ decreased beginning at D08 (*p* = 0.006). The cADPR/NAD⁺ ratio did not differ significantly across ages (*p* = 0.189). These ratios reflect relative metabolite abundances and may be influenced by production, recycling, and clearance; they do not directly measure NAD⁺ consumption or enzyme-specific NADase activity. **(D)** Integrated NAD⁺ network map showing D15 changes relative to D03. Metabolite node colors indicate log₂ fold changes: blue, decreased abundance; rose, increased abundance. Thick node outlines indicate FDR-adjusted *q* < 0.05; thin outlines indicate *q* ≥ 0.05. Boxed enzyme labels show protein log₂ fold changes, with thick outlines identifying significant changes. Thin gray boxes identify enzymes not detected in the proteomic dataset. The map highlights reductions in several *de novo* pathway components alongside positive fold changes in Preiss–Handler-associated metabolites, downstream cofactors, and methylated pyridine metabolites. Metabolite accumulation does not independently establish increased pathway flux or precursor disposal. **(E)** Comparison of enzyme log₂ fold changes with associated metabolite-ratio changes at D08, D12, and D15. Marker shapes indicate age: diamonds, D08; squares, D12; triangles, D15. Filled markers indicate that both the protein and metabolite-ratio changes met the stated significance criterion (*q* < 0.05); open markers indicate that at least one did not. Blue and orange identify concordant and discordant directional changes, respectively. The dashed diagonal indicates the 1:1 line. Metabolite ratios are interpreted as pathway indices rather than direct measurements of enzymatic activity or flux. **(F)** Temporal changes in respiratory-chain proteins and associated metabolite ratios. Upper gauges show the proportion of detected subunits in each respiratory complex that differed significantly from D03 (*q* < 0.05). Dashed lines indicate the age-matched proportion of altered proteins among all detected metabolic proteins: 17% at D08, 35% at D12, and 62% at D15. Gray and rose fills distinguish the background component and any excess above that background. At D08, 43% of detected Complex I subunits were altered, whereas none of the detected Complex III, IV, or V subunits met the significance criterion. At D12, the proportions were 65% for Complex I and 42% for Complex III. At D15, they were 83%, 75%, 86%, and 50% for Complexes I, III, IV, and V, respectively. Complex II was insufficiently detected and is designated not testable (n.t.). Gauge widths are proportional to the numbers of detected subunits. The lower portion of (F) shows hypoxanthine/ADP, coenzyme A (CoA)/pantothenate, flavin adenine dinucleotide (FAD)/riboflavin, GSSG/GSH, aspartate/glutamine, and fumarate/α-ketoglutarate ratios as log₂ fold changes relative to D03. Filled bars indicate nominal *p* < 0.05 versus D03; open bars indicate *p* ≥ 0.05. Ratios were measured in whole-worm extracts. Their placement within the mitochondrial schematic denotes associated biochemical processes rather than experimentally established subcellular localization. Analyses used three independent biological replicates per age. Overall age effects in (B) and (C) were assessed by one-way ANOVA; reported *p* values are unadjusted. Protein and metabolite *q* values in (D–F) were calculated using Benjamini–Hochberg FDR correction. Metabolite-ratio comparisons in the lower portion of (F) used unpaired, two-tailed Welch’s *t* tests against D03 and were not FDR-adjusted. Significance is encoded graphically as described above.

Among the proteomic functional groups examined, Complex I showed an early and sustained decline (**Figure 6B**). The mean abundance of 23 detected subunits decreased sharply between D03 and D08 and continued to decline through D15 (overall age effect, *p* < 0.001). Many individual subunits were significantly reduced at D08, and most were reduced by D15. GAS-1 was the only detected Complex I subunit to show a significant increase (**Figure S6A**).

Other functional groups changed later. NAD-dependent dehydrogenases, NADPH-producing enzymes, and nucleotidases remained near or above their D03 values at D08 before declining at subsequent ages. None of the 23 NAD-dependent dehydrogenases differed significantly at D08, and most significant reductions emerged only at D15. Among NADPH-producing enzymes and nucleotidases, IDH-1, NTFC-1, and BPNT-1 decreased significantly at D12 and D15 (**Figure S6A**). Thus, broad Complex I protein loss preceded the reductions observed in several other NAD⁺-associated functional groups.

Consumption-associated metabolite ratios also followed distinct trajectories (**Figure 6C**). NAM/NAD⁺ declined sharply at D08 and remained low through D15 (overall age effect, *p* = 0.006), whereas cADPR/NAD⁺ did not differ significantly across ages (*p* = 0.189). The lower NAM/NAD⁺ ratio could reflect the expanding NAD⁺ pool, altered NAM production, or changes in NAM recycling and clearance. These ratios therefore describe relationships among metabolite pools rather than directly measuring NAD⁺ consumption or NADase activity.

Mapping D15 changes onto the NAD⁺ network further illustrated the divergence among pathway components (**Figure 6D**). NFK, Kyn, and kynurenate (kynurenic acid) were lower than at D03, alongside reduced TDO-2 and NKAT-3 abundance. In contrast, NA, NaMN, NAD⁺, and NADP⁺ showed positive fold changes. Trigonelline and 1-methylnicotinamide also increased, consistent with changes in methylated pyridine metabolite pools (see also **Figure S5E**). NAM methylation to 1-methylnicotinamide has been linked to sirtuin-dependent lifespan extension in *C. elegans* **[79]**, making this branch relevant to the physiological interpretation of network remodeling. However, accumulation alone cannot distinguish increased formation from reduced utilization or clearance.

To assess correspondence between protein and metabolite changes, we compared enzyme fold changes with associated product-to-substrate ratios (**Figures 6E** and **S6B**). Only comparisons involving TDO-2 showed significant concordant reductions in both measures, with lower TDO-2 abundance accompanied by reduced NFK/Trp and Kyn/Trp ratios at all three later ages. The NKAT-3-associated KynA/Kyn ratio also fell within the concordant-decrease quadrant, but both measures did not jointly meet the significance criteria. Comparisons involving FLU-2, HAAO-1, and NPRT-1 were frequently discordant. Most notably, NaMN/NA decreased at every later age despite unchanged or slightly increased NPRT-1 abundance (**Figure S6B**). These patterns indicate that protein abundance alone does not explain the metabolite trajectories and suggest additional contributions from precursor availability, pathway regulation, consumption, and clearance.

Analysis across the electron transport chain reinforced the early emergence of Complex I remodeling (**Figure 6F**). At D08, 43% of detected Complex I subunits met the threshold for change relative to D03, compared with 17% of all detected metabolic proteins. No detected subunits of Complexes III, IV, or V met this threshold at D08. By D12, the proportions increased to 65% for Complex I and 42% for Complex III, while Complexes IV and V remained comparatively preserved. At D15, changes affected 83% of detected Complex I subunits, 75% of Complex III, 86% of Complex IV, and 50% of Complex V. Complex II was insufficiently detected for evaluation.

At every sampled age, the proportion of altered Complex I subunits exceeded the background proportion among all detected metabolic proteins. In contrast, the proportion of altered TCA cycle enzymes remained below this background through D12, while tryptophan-degradation enzymes closely tracked it (**Figure S6D**). Among the respiratory complexes assessed, Complex I therefore exhibited the earliest broad protein-abundance changes, followed by more extensive respiratory-chain remodeling later in adulthood.

Together, these analyses place early Complex I remodeling alongside divergent NAD⁺ pathway metabolite profiles, reduced mid-adult precursor incorporation, and altered redox and energetic status. Complex I changes preceded broad remodeling of the other measured respiratory complexes, but their contribution to the accompanying metabolic transition remains unresolved. Direct measurements of respiratory activity will be needed to establish their functional consequences. We next examined the physiological importance of NAD⁺-linked pathways by testing how consumer and salvage mutations, together with precursor supplementation, affected locomotion, reproduction, and lifespan.

### Disruption of NAD⁺-consuming and salvage enzymes compromises fitness, while high-dose NAM and trigonelline shorten lifespan

The maintained or increased expression of selected NAD⁺ consumer genes during adulthood prompted us to examine their physiological importance. We measured locomotion, reproduction, and lifespan in animals carrying loss-of-function mutations in the NAD⁺ consumers *parp-1*, *parp-2*, *tank-1*, and *tir-1*, or the nicotinamide salvage enzyme *pnc-1*.

Locomotor phenotypes differed among genotypes and assay conditions (**Figure 7A** and **7B**). Crawling speed on solid media was significantly lower in *parp-1*, *parp-2*, and *tir-1* mutants than in wild-type N2 animals, whereas *tank-1* and *pnc-1* mutants did not differ significantly from wild type (**Figure 7A**). In liquid media, thrashing rates decreased in *parp-2* and *pnc-1* mutants but increased modestly in *tank-1* mutants. Neither *parp-1* nor *tir-1* mutants differed significantly from wild type in this assay (**Figure 7B**). These responses indicate distinct contributions of individual NAD⁺-linked enzymes to locomotor performance.

**Figure 7.**
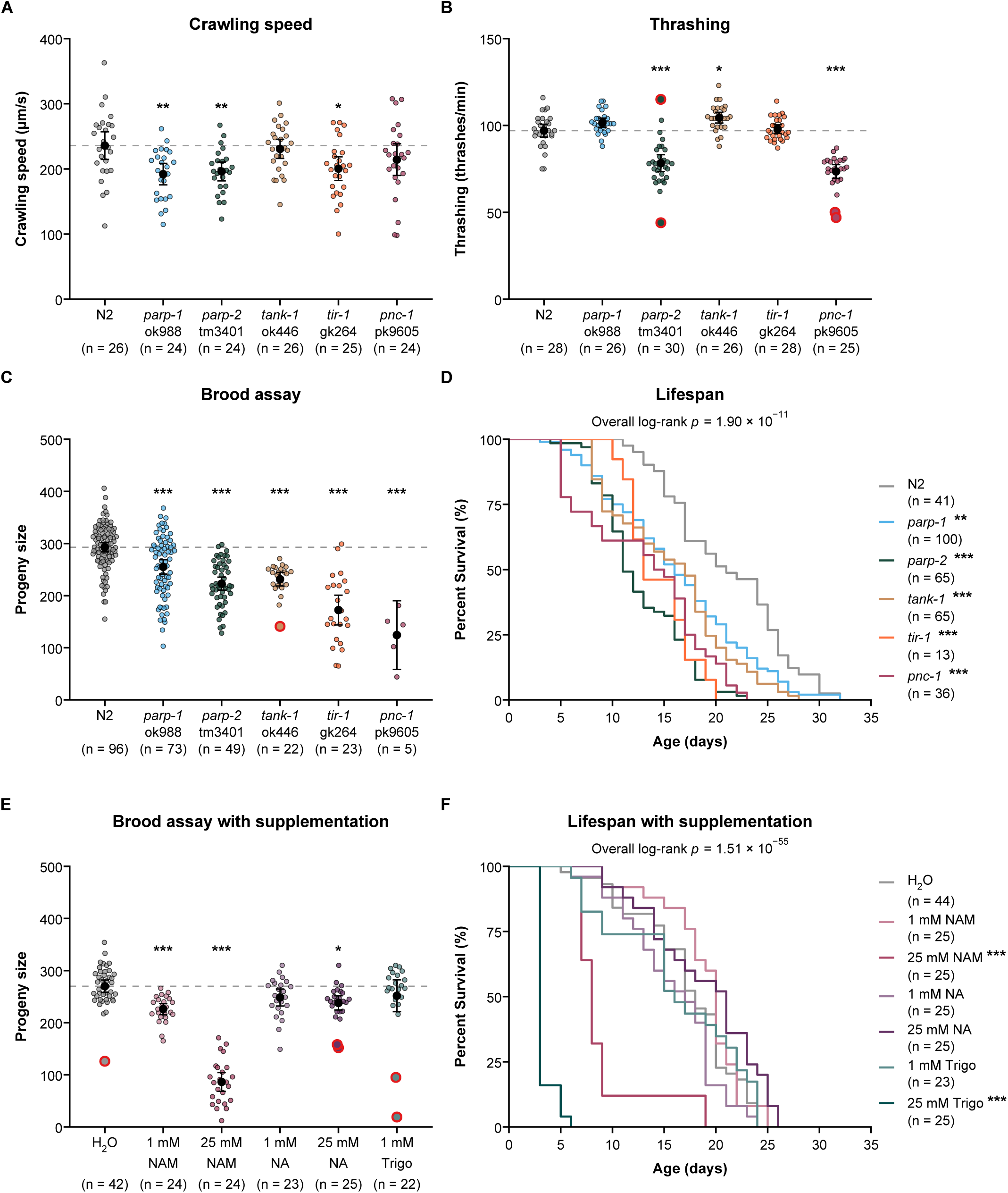
| Disruption of NAD⁺-consuming and salvage enzymes compromises fitness, while high-dose NAM and trigonelline shorten lifespan. **(A)** Crawling speed on unseeded nematode growth medium in wild-type N2 animals and loss-of-function mutants of the NAD⁺ consumers *parp-1, parp-2, tank-1,* and *tir-1*, or the nicotinamide salvage enzyme *pnc-1*. Crawling speed was significantly lower in *parp-1, parp-2,* and *tir-1* mutants than in wild type. Neither *tank-1* nor *pnc-1* mutants differed significantly from wild type. **(B)** Liquid thrashing rates in the same genotypes. Rates decreased in *parp-2* and *pnc-1* mutants and increased modestly in *tank-1* mutants. Neither *parp-1* nor *tir-1* mutants differed significantly from wild type. **(C)** Total brood size, measured as viable progeny per animal. All five mutant strains produced significantly fewer progeny than wild type. The smaller evaluable *pnc-1* cohort reflected internal hatching (“bagging”) associated with its reproductive defects. **(D)** Kaplan–Meier survival curves for wild-type and mutant animals under standard culture conditions. All five mutant strains had significantly shorter lifespans than wild type (overall log-rank *p* = 1.9 × 10⁻¹¹). Adjusted pairwise significance relative to wild type is indicated beside each genotype. **(E)** Total brood size of wild-type animals supplemented with nicotinamide (NAM; 1 or 25 mM), nicotinic acid (NA; 1 or 25 mM), or trigonelline (1 mM), relative to the H₂O vehicle control. Both NAM concentrations reduced brood size, with the largest reduction at 25 mM. Supplementation with 25 mM NA also reduced brood size, whereas 1 mM NA and 1 mM trigonelline did not differ significantly from the control. Brood size was not assessed following exposure to 25 mM trigonelline. **(F)** Kaplan–Meier survival curves for wild-type animals receiving H₂O vehicle, NAM (1 or 25 mM), NA (1 or 25 mM), or trigonelline (1 or 25 mM). The overall treatment effect was significant (log-rank *p* = 1.5 × 10⁻⁵⁵). Exposure to 25 mM trigonelline resulted in near-complete mortality by Days 5–6, while approximately 12% of animals receiving 25 mM NAM survived to Day 11. Survival with 1 mM NAM, 1 or 25 mM NA, or 1 mM trigonelline did not differ significantly from the vehicle control. For (A–C) and (E), colored points represent individual animals; black circles and error bars indicate means and 95% confidence intervals. Red rings identify observations flagged by Rosner’s generalized extreme Studentized deviate test. All flagged observations were retained in the analyses. Exact group sizes are displayed beneath each group and represent animals scored across independent experiments. Group-size ranges are: (A), *n* = 24–26; (B), *n* = 25–30; (C), *n* = 5–96; (D), *n* = 13–100; (E), *n* = 22–42; and (F), *n* = 23–44. Outcomes in (A–C) and (E) were analyzed by one-way ANOVA followed by Dunnett’s multiple-comparison test against wild type (A–C) or vehicle control (E). Survival curves in (D) and (F) were compared using overall log-rank (Mantel–Cox) tests followed by pairwise log-rank tests with Holm correction. *p < 0.05, **p < 0.01, and ***p < 0.001; ns, not significant.

Reproductive defects were more widespread. All five mutant strains produced significantly fewer progeny than wild-type animals, with the largest reductions in *tir-1* and *pnc-1* mutants and intermediate reductions in *parp-1*, *parp-2*, and *tank-1*mutants (**Figure 7C**). The smaller evaluable *pnc-1* cohort reflected internal hatching, or “bagging,” associated with its reproductive defects **[63, 68, 80]**. Thus, disruption of either NAD⁺ consumers or nicotinamide salvage reduced reproductive output under the conditions tested.

All five mutant strains also exhibited significantly shorter lifespans than wild type (**Figures 7D** and **S7A–S7E**). Mean lifespan decreased by 23.3% in *parp-1*, 25.7% in *tank-1*, 31.4% in *tir-1*, 38.0% in *pnc-1*, and 39.9% in *parp-2* mutants. The *pnc-1* phenotype was consistent with the reduced lifespan previously reported following *pnc-1* knockdown **[81]**. These findings support physiological roles for NAD⁺-consuming and salvage enzymes in maintaining survival under standard culture conditions. Because the mutations were present throughout development, however, the observed phenotypes cannot distinguish developmental contributions from functions specific to adult aging.

We next tested whether supplementation with NAD⁺ precursors or related pyridine metabolites altered reproduction and lifespan in wild-type animals (**Figures 7E and 7F**). NAM reduced total brood size at both 1 mM and 25 mM, with a larger reduction at 25 mM (**Figure 7E**). Supplementation with 25 mM NA modestly reduced brood size, whereas 1 mM NA and 1 mM trigonelline had no significant effect. Animals exposed to 25 mM trigonelline laid eggs but died before their total brood size could be determined.

Survival outcomes likewise depended on metabolite identity and concentration (**Figures 7F** and **S7F–S7K**). Neither 1 mM NAM, 1 mM NA, 25 mM NA, nor 1 mM trigonelline significantly altered lifespan relative to the water vehicle control; mean lifespan changes ranged from −7.6% to +9.1% (**Figure S7F** and **S7H–S7J**). In contrast, 25 mM NAM reduced mean lifespan by 47.5%, with approximately 12% of animals surviving to Day 11 (**Figure S7G**), consistent with prior reports of reduced survival on NAM-containing medium **[81]**. Exposure to 25 mM trigonelline produced an earlier and more severe response: nearly all animals died by Days 5–6, and mean lifespan decreased by 80.4% (**Figure S7K**). Water-treated controls from two independent experiments showed no significant difference in survival, with a 3.0% difference in mean lifespan, and were combined for the displayed treatment comparisons (**Figure S7L**). None of the tested supplementation regimens significantly extended lifespan.

Together, these experiments reveal gene-specific locomotor phenotypes alongside broadly reduced reproduction and survival following disruption of NAD⁺-consuming or salvage enzymes. Supplementation effects also differed by endpoint: 1 mM NAM reduced brood size without significantly altering lifespan, whereas high-dose NAM and trigonelline markedly reduced survival. These findings support the physiological importance of NAD⁺-linked pathways and show that increasing precursor availability alone does not necessarily improve fitness or longevity.

## Discussion

Age-associated changes in NAD⁺ abundance have been linked to metabolic and mitochondrial dysfunction, but pool size alone provides an incomplete view of NAD⁺ homeostasis **[20, 25, 28, 78]**. Steady-state measurements cannot distinguish changes in synthesis, consumption, recycling, or clearance, allowing substantial network remodeling to occur without a corresponding decline in NAD⁺ abundance **[20, 25, 78]**. Here, we integrated LC-MS metabolomics, proteomics, gene-expression profiling, and *in vivo* D₄-NAM and D₄-NA tracing **[78]** across the *C. elegans* adult lifespan. NAD⁺ accumulated under standard plate conditions despite reductions in selected biosynthetic transcripts and proteins. This accumulation coexisted with reduced mid-adult precursor incorporation, pronounced changes in NR-associated metabolism, and persistent alterations in redox and energetic indices. Together, these findings show that preservation of the NAD⁺ pool does not necessarily indicate preservation of the metabolic processes that sustain and use it.

The integrated datasets revealed selective remodeling rather than a uniform decline in NAD⁺ biosynthesis. Reduced *afmd-1* expression, numerical decreases in *tdo-2* transcripts, lower TDO-2 and UMPS-1 protein abundance, and declining levels of selected kynurenine-pathway metabolites contrasted with maintained or increased Preiss–Handler-associated components. Increased *nprt-1* and *qns-1* expression further distinguished the transcriptional responses within these branches. NAM recycling through PNC-1 connects precursor salvage to the Preiss–Handler pathway **[63, 68]**, and incorporation of both NAM– and NA-derived label into NAD⁺ demonstrated continued precursor utilization across adulthood. However, without direct tracing of tryptophan-derived synthesis, these experiments cannot quantify the relative contributions of *de novo* synthesis and salvage or establish compensatory transfer between them. The findings instead reveal that NAD⁺ abundance can be maintained alongside substantial reorganization of its supporting network, consistent with the broader concept of metabolic adaptation during physiological stress **[82]**.

Tracing exposed changes that were not apparent from steady-state NAD⁺ abundance. NAD⁺ labeling from both precursors decreased at D08 and D12 before partially recovering at D15. This pattern complements observations in aged mice, where NAD⁺ synthesis flux is maintained despite lower tissue concentrations **[25]**. In both settings, pool size and precursor dynamics provide distinct information about NAD⁺ metabolism. The isotope distributions also connect precursor incorporation with downstream processing: loss of deuterium during NAD(H) redox cycling generates M+3 NAD⁺, and subsequent consumption can release labeled NAM for recycling through biosynthetic pathways. Nevertheless, the four-hour endpoint does not independently resolve absolute synthesis, consumption, or recycling rates. Precursor uptake, intracellular enrichment, pool size, and redox cycling can each influence the observed labeling.

NR-associated metabolism emerged as a particularly age-sensitive feature. Steady-state NR abundance declined significantly, while NA-derived NR labeling became undetectable at D08, D12, and D15. NAM-derived NR labeling also decreased significantly at all three ages, although low residual labeling remained at D08 and D15. Reduced NMN labeling under both tracer conditions placed these changes within broader remodeling of precursor handling. Importantly, these findings do not establish loss of NR utilization capacity or predict the response to exogenous NR. Mammalian studies illustrate that NR handling can involve both kinase-mediated salvage and extracellular hydrolysis **[39, 55]**, underscoring the need to distinguish precursor formation from utilization. Direct NR tracing, coupled with perturbation of relevant salvage and recycling enzymes, will help determine which processes account for the age-dependent loss of label accumulation.

The contrasting NAD⁺ trajectories across culture conditions raise a related question about physiological demand. NAD⁺ accumulated across adulthood on NGM plates but increased at D08 and declined numerically thereafter during liquid incubation with continuous rocking. Although the latter differences were not significant, the pattern occurred with both tracers. Sustained swimming may impose greater energetic demand than plate culture, where animals can move intermittently and potentially conserve energy. However, the experiments also differed in environmental conditions and precursor exposure. The results therefore motivate, rather than establish, the hypothesis that limitations in NAD⁺ maintenance become more apparent under increased demand. Matched experiments that manipulate movement while controlling culture conditions will be needed to test this possibility.

Extracellular labeling and methylated metabolite profiles extend the remodeling beyond intracellular NAD⁺ abundance. Following D₄-NAM exposure, labeled NA represented approximately 30–40% of the extracellular NA pool, whereas labeled NAM represented less than 10% of the extracellular NAM pool following D₄-NA exposure. These asymmetric patterns are consistent with NAM deamination and extracellular appearance of labeled NA, highlighting precursor exchange as a potential component of network organization. Related exchange processes contribute to mammalian host–microbiome NAD⁺ metabolism **[83]**. However, extracellular enrichment does not quantify the fraction of administered precursor exported or establish net precursor loss. Likewise, methylated metabolite accumulation should not automatically be interpreted as disposal: 1-methylnicotinamide has been linked to longevity in *C. elegans* **[79]**, and trigonelline can support NAD⁺ metabolism in other settings **[76]**. Changes in their abundance may reflect altered formation, utilization, clearance, or signaling functions.

Network remodeling coincided with distinct redox and energetic trajectories. Adenylate indices shifted beginning at D08, while GSH/GSSG and lactate/pyruvate showed sustained reductions that became significant later in adulthood. NAD⁺/NADH increased numerically without reaching significance, and the mid-adult elevation in ox-NAD/NAD⁺ subsided by D15. Changes in SAM/SAH, MeNAM/NAM, carnitine-associated ratios, and TCA metabolite ratios further placed NAD⁺ remodeling within a broader metabolic transition. These measurements extend the relationship between NAD⁺ metabolism and mitochondrial physiology **[28]**, but their interpretation remains constrained by whole-animal sampling and semi-quantitative peak-area measurements. They do not establish compartment-specific redox states, absolute molar energy charge, or changes in individual reaction fluxes.

Complex I showed the earliest broad proteomic remodeling among the respiratory complexes assessed. At D08, 43% of detected Complex I subunits were altered, before comparable changes emerged in Complexes III, IV, and V. The proportion of altered Complex I proteins exceeded the background proportion among detected metabolic proteins at every sampled age. This temporal pattern identifies Complex I as a candidate contributor to the accompanying redox and energetic changes. However, reduced subunit abundance does not directly establish impaired electron transport, and the shared timing cannot determine whether Complex I remodeling initiates or responds to the metabolic transition. Respiratory measurements and adult-onset interventions will be essential to resolve this relationship.

The genetic experiments demonstrate that NAD⁺-linked enzymes contribute to organismal fitness, although their effects differ across physiological endpoints. All five consumer or salvage mutants showed reduced reproductive output and shorter lifespan, whereas locomotor phenotypes varied by genotype and assay. These results emphasize that benefits reported for consumer inhibition in other settings cannot be assumed to apply to lifelong loss of function **[37]**. Pharmacological PARP inhibition has been reported to increase NAD⁺ and extend lifespan in *C. elegans* **[18]**. The different outcomes may reflect intervention timing, degree of inhibition, genetic background, culture conditions, or pharmacological effects beyond the intended target. Developmental functions and roles beyond NAD⁺ consumption may also contribute to the mutant phenotypes observed here.

Supplementation similarly produced metabolite-, dose-, and endpoint-dependent outcomes. None of the tested regimens significantly extended lifespan. NAM at 1 mM reduced brood size without significantly altering survival, whereas 25 mM NAM and trigonelline markedly shortened lifespan. Previous studies have reported both reduced survival on NAM-containing medium **[81]** and NAM-associated lifespan extension involving methylation **[79]**. Together, these observations reinforce the importance of dose, metabolic context, and the physiological outcome being measured when evaluating NAD⁺-directed interventions **[42, 63, 81]**. Our experiments do not establish that the adverse supplementation outcomes resulted from increased NAD⁺ abundance or a specific downstream mechanism.

Several questions follow from these findings. First, the temporal association between the mid-adult transition and declining reproduction does not distinguish reproductive influences from chronological aging. Comparing fertile animals with carefully controlled reproductive perturbations could help separate germline-dependent effects from somatic metabolic changes, while accounting for the metabolic consequences of sterility and temperature shifts. Second, whole-animal profiles may conceal opposing responses among tissues. Adult-onset, tissue-specific perturbations paired with metabolic measurements could identify where precursor recycling, NAD⁺ consumption, and mitochondrial remodeling contribute most strongly to the observed phenotypes. Finally, compartment-targeted NAD(H) and ATP biosensors could resolve temporal and subcellular changes that are obscured by pooled extracts. Combined with respiratory measurements and mitochondrial imaging, these approaches would connect network remodeling to its anatomical and functional consequences.

Overall, aging in *C. elegans* involves extensive NAD⁺ network remodeling despite maintained or increased steady-state NAD⁺ abundance. Reduced precursor incorporation, altered NR-associated metabolism, early Complex I protein changes, and persistent shifts in redox and energetic indices reveal metabolic features that would be missed by measuring NAD⁺ alone. Defining how these processes interact will help distinguish interventions that increase the NAD⁺ pool from those that preserve the physiological functions it supports.

## RESOURCE AVAILABILITY

### Lead contact

Further information and requests for resources and reagents should be directed to the lead contact Melanie McReynolds.

### Materials availability

The study did not generate new materials.

### Data and code availability

- All metabolomics, proteomics, and isotope tracing data reported have been deposited at MassIVE (Mass Spectrometry Interactive Virtual Environment) and are publicly available as of the date of publication.
- Custom R scripts and computational workflows used for metabolomics and proteomics downstream analysis are available upon request.
- Any additional information required to reanalyze the data reported in this paper is available from the lead contact upon request.

## Acknowledgments

We would like to thank Professor Wendy Hanna-Rose for her thoughtful feedback and suggestions for the manuscript. We would like to acknowledge the Huck Institutes’ Metabolomics Core Facility (RRID:SCR_023864) for use of the OE 240 LC-MS and Sergei Koshkin for helpful discussions on sample preparation and analysis. We also acknowledge the Proteomics and Mass Spectrometry Core Facility at the Huck Institutes of the Life Sciences, Pennsylvania State University, for use of the Thermo Eclipse Tribrid Mass Spectrometer, technical assistance, and Tatiana N. Laremore for helpful discussions on sample preparation and analysis (RRID:SCR_024462). We would like to thank Alexandria Claire Murphy and Julia Mudryk for taking care of the LCMS and the supplies respectively. This work was funded and supported by the Howard Hughes Medical Institute (HHMI) Hanna H. Gray Fellows Program Faculty Phase (Grant# GT15655, M.R.M), the Burroughs Welcome Fund PDEP Transition to Faculty (Grant# 1022604, M.R.M), and the National Science Foundation (NSF) Faculty Early Career Development (CAREER) Program (Grant #2441332, M.R.M.).

## Author contributions

M.R.M. and A.A.A. designed the study. A.A.A., M.S.Y., I.A.A., A.M.S., J.J.S., S.V.J., I.J.M., B.C.B., K.N.D., E.O., E.R.R. performed experiments and analyzed the data. M.R.M. and A.A.A. planned the project and supervised the experiments. A.A.A., M.S.Y., and T.N.L. led the proteomics experiment and analysis. M.R.M., A.A.A. wrote the initial manuscript. A.A.A., M.S.Y., I.A.A., A.M.S., J.J.S., S.V.J., I.J.M., B.C.B., K.N.D., E.O., E.R.R., T.N.L., and M.R.M. edited and approved the manuscript.

## Declaration of generative AI use

During the preparation of this work, the authors used ChatGPT (OpenAI) to assist with improving readability, grammar, and clarity of the text, and Claude (Anthropic) to assist with coding for data visualization. After using these tools, the authors reviewed and edited the content as needed and take full responsibility for the content of the published article.

## STAR METHODS

Detailed methods are provided in the online version of this paper and include the following:

- KEY RESOURCES TABLE
- RESOURCE AVAILABILITY
  ○ Lead contact
  ○ Materials availability
  ○ Data and code availability
- EXPERIMENTAL MODEL AND SUBJECT DETAIL
  ○ *C. elegans* culture and strains
- METHODS DETAILS
  ○ Lifespan Analysis
  ○ Brood–Reproductive Assay
  ○ Crawling-speed Analysis (WormLab)
  ○ Thrashing Assay
  ○ Sample collection and metabolite extraction
  ○ Sample preparation for LCMS
  ○ Stable-isotope tracing
  ○ Metabolite measurement LCMS
  ○ RNA isolation and qRT-PCR
  ○ Proteomics Sample preparation and peptide purification
  ○ Proteomics Workflow
  ○ Precursor Supplementation
  ○ Integrative multi-omics network analysis (need to write—Kareem/Mo!)
- QUANTIFICATION AND STATISTICAL ANALYSIS
  ○ Statistical analysis

## SUPPLEMENTAL INFORMATION

Supplemental information can be found online.

## Abbreviations

3-HAA: 3-hydroxyanthranilic acid
ACMS: 2-amino-3-carboxymuconate semialdehyde
ACN: acetonitrile
ADP: adenosine diphosphate
AGC: automatic gain control
AHPDB: 4-(2-amino-3-hydroxyphenyl)-2,4-dioxobutanoate
AMP: adenosine monophosphate
ANOVA: analysis of variance
ATP: adenosine triphosphate
BCA: bicinchoninic acid
cADPR: cyclic ADP-ribose
CD157: cluster of differentiation 157 (bone marrow stromal cell antigen 1, BST1)
CD38: cluster of differentiation 38
cDNA: complementary DNA
CGC: Caenorhabditis Genetics Center
CoA: coenzyme A
Ct: cycle threshold
D03: 62 h after egg laying
D08, D12, D15: Days 8, 12, and 15
D₄-NA: labeled nicotinic acid
D₄-NAM: labeled nicotinamide
FAD: flavin adenine dinucleotide
FDR: false discovery rate
FUdR: 5-fluoro-2′-deoxyuridine
GSH: reduced glutathione
GSSG: glutathione disulfide (oxidized glutathione)
H₂O: water
HILIC: hydrophilic interaction liquid chromatography
HSD: honestly significant difference
IQR: interquartile range
L1, L3, L4: first, third, and fourth larval stages
LC-MS: liquid chromatography–mass spectrometry
LC-MS/MS: liquid chromatography–tandem mass spectrometry
LDH: lactate dehydrogenase
M+n: mass isotopologue containing n deuterium atoms (M+0, unlabeled)
MassIVE: Mass Spectrometry Interactive Virtual Environment
MeOH: methanol
n.t.: not testable
NA: nicotinic acid
NAD⁺: nicotinamide adenine dinucleotide (oxidized)
NADH: nicotinamide adenine dinucleotide (reduced)
NADP⁺: nicotinamide adenine dinucleotide phosphate (oxidized)
NADPH: nicotinamide adenine dinucleotide phosphate (reduced)
NAM: nicotinamide
NaMN: nicotinic acid mononucleotide
NAMPT: nicotinamide phosphoribosyltransferase
NGM: nematode growth medium
NMN: nicotinamide mononucleotide
NR: nicotinamide riboside
ns: not significant
ox-NAD: Pyridone-containing adenine dinucleotides
PARP: poly(ADP-ribose) polymerase
PC1, PC2: first and second principal components
PCR: polymerase chain reaction
PLS-DA: partial least-squares discriminant analysis
Q²: cross-validated predictive ability
QA: quinolinate
qRT-PCR: quantitative reverse-transcription PCR
R²Y: explained variance of the class response (goodness of fit)
RRID: Research Resource Identifier
SARM1: sterile alpha and Toll/interleukin-1 receptor motif-containing protein 1
SD: standard deviation
SEM: standard error of the mean
TIR: Toll/interleukin-1 receptor
UV: ultraviolet
WHR: Wendy Hanna-Rose laboratory

## Supplementary Figure Legends

**Figure S1| Pathway-level and replicate-level analysis of NAD⁺-related transcripts.**

(**S1A**) Heatmap of mean log₂ fold changes relative to D03 for the 22 NAD⁺-related genes in **Figure 1C** at D08, D12, and D15, grouped by pathway: de novo, Preiss–Handler, NR kinase (NRK), salvage, sirtuins, PARPs/TIR, and NADP⁺ synthesis. (**S1B**) Log₂ fold changes relative to D03 for each gene. Bars show mean ± SEM; points represent individual biological replicates. Expression was measured by qRT-PCR and normalized to the geometric mean of *cdc-42*, *cyc-1*, and *pmp-3*. Data represents six independent biological replicates per age. Statistical significance was determined by one-way ANOVA followed by Dunnett’s multiple-comparison test against D03. *p < 0.05, **p < 0.01, and ***p < 0.001.

**Figure S2| Progressive decline of the NAD⁺-associated proteome across adulthood.**

(**S2A and S2B**) Volcano plots of protein-abundance changes at D08 (**S2A**) and D12 (**S2B**) relative to D03 for 124 NAD⁺-related and NAD⁺-dependent proteins. Vertical dashed lines indicate |log₂ fold change| = 1. Twenty-three and 41 proteins decreased significantly at D08 and D12, respectively, and one and two proteins increased. Labeled NAD⁺ biosynthetic enzymes are shown as filled (both criteria met) or open (criteria not met) circles. (**S2C**) Log₂ fold changes relative to D03 for TDO-2, UMPS-1, NKAT-3, FLU-2, HAAO-1, and NPRT-1. Bars show mean ± SD; points represent individual biological replicates. (**S2D**) Log₂ fold changes relative to D03 for the 124 proteins grouped into 12 functional categories. Numbers in parentheses indicate proteins per category, and vertical black lines indicate category medians. Blue and red points indicate significantly decreased and increased proteins, respectively; gray points indicate proteins that did not meet both criteria. Data represent three independent biological replicates per age. Statistical significance was assessed using moderated two-sided t tests with Benjamini–Hochberg FDR correction across all 1,810 quantified proteins. Proteins were classified as altered when q < 0.05 and |log₂ fold change| > 1; asterisks in (**S2C**) indicate that both criteria were met.

**Figure S3| Extended profiling of the steady-state NAD⁺ metabolome.**

(**S3A**) Variable importance in projection (VIP) scores from the PLS-DA model in **Figure 3A**. Bar colors indicate pathway assignment as in (B); the dashed line marks VIP = 1. (**S3B**) Heatmap of mean log₂ fold changes relative to D03 for all 24 measured metabolites at D08, D12, and D15, grouped by pathway assignment. (**S3C–K**) Normalized abundance of N-formylkynurenine (NFK; **S3C**), 3-hydroxykynurenine (3-HK; **S3D**), kynurenic acid (**S3E**), 4-(2-Amino-3-hydroxyphenyl)-2,4-dioxobutanoate (AHPDB; **S3F**), 2-aminomuconate semialdehyde (2-AMS; **S3G**), 2-aminomuconate (**S3H**), picolinic acid (**S3I**), nicotinic acid riboside (NaR; **S3J**), and cyclic ADP-ribose (cADPR; **S3K**). Metabolite signals were normalized to total ion count and BCA-measured protein abundance. Boxplots show medians and IQRs, with whiskers extending to 1.5 × IQR. Large, outlined symbols represent the mean of technical injections for each biological replicate; smaller translucent symbols represent individual injections. Data represent four independent biological replicates per age. Statistical significance was determined by one-way ANOVA followed by Tukey’s HSD test; asterisks in (**S3B–K**) indicate significant differences from D03. *p < 0.05, **p < 0.01, and ***p < 0.001.

**Figure S4| Fate of the deuterium label from D₄-NAM and D₄-NA in the NAD⁺ network**.

(**S4A**) D₄-NAM and D₄-NA carry deuterium (²H) at C2, C4, C5, and C6 of the pyridine ring. D₄-NAM is deamidated to D₄-NA by PNC-1 and PNC-2, and both tracers enter NAD⁺ through the Preiss–Handler pathway (NPRT-1, NMAT-1/NMAT-2, and QNS-1), generating M+4 NAD⁺. Reduction of NAD⁺ to NADH adds an unlabeled hydride at C4. Re-oxidation removes either the unlabeled hydrogen, regenerating M+4 NAD⁺, or the C4 deuterium, generating M+3 NAD⁺. NAD⁺ consumers release M+3 NAM, which can be deamidated to M+3 NA. M+3 NMN arises from NAD⁺ turnover and is converted to NAD⁺ by NMAT-1/NMAT-2; NR enters through NMRK-1. Solid arrows indicate synthesis, dashed arrows indicate consumption, and double arrows indicate tracer uptake. R, ADP-ribose.

**Figure S5| Additional metabolite ratios reporting on methylation, carnitine, and TCA cycle status.**

(**S5A–F**) Peak-area ratios of S-adenosylmethionine/S-adenosylhomocysteine (SAM/SAH; **S5A**), acetylcarnitine/free carnitine (C2/C0; **S5B**), α-ketoglutarate/succinate (**S5C**), succinate/fumarate (**S5D**), N¹-methylnicotinamide/nicotinamide (MeNAM/NAM; **S5E**), and glutamine/glutamate (**S5F**). (**S5G and S5H**) Quality-control ratios for ATP breakdown (**S5G**) and adenylate kinase (AK) equilibrium (**S5H**). (**S5I–S5L**) Fold changes of the individual metabolites contributing to the ratios in (**S5A–S5D**), normalized to the D03 mean and displayed on a log₂ scale. Black symbols and lines show mean ± SEM; gray symbols in (**S5A–S5H**) represent individual biological replicates. Data represent three independent biological replicates per age. Values are semi-quantitative LC-MS peak-area ratios from whole-animal extracts. Statistical significance was determined by one-way ANOVA of log₂-transformed ratios followed by Tukey’s HSD test; asterisks indicate significant differences from D03. *p < 0.05, **p < 0.01, and ***p < 0.001.

**Figure S6| Protein-level and pathway-level details of the multi-omic integration**.

**S6A**) Heatmap of log₂ fold changes relative to D03 for NAD⁺-relevant proteins at D08, D12, and D15, grouped into de novo, Preiss–Handler, Complex I (23 proteins), NAD⁺-dependent dehydrogenase (23), NADPH-producing (4), and nucleotidase (3) modules. (**S6B**) Labeled version of **Figure 6E** comparing enzyme log₂ fold changes with changes in the corresponding product-to-substrate metabolite ratios. Each point is labeled with its enzyme and ratio; colors indicate age. Filled circles indicate that both changes were significant; open circles indicate that at least one was not. Shaded quadrants indicate concordant changes, and the dashed diagonal indicates the 1:1 line. (**S6C**) Trajectories of individual metabolites in the de novo arm (Trp, NFK, Kyn, and 3-HK) and Preiss–Handler arm (NA, NaMN, NaR, and NAD⁺) summarized in **Figure 6A**. Open symbols show member means; filled symbols and shading show the arm mean ± SEM across metabolites. (**S6D**) Percentage of detected proteins differing significantly from D03 in Complex I, Complex III, Complex IV, the TCA cycle, and tryptophan degradation. Numbers in parentheses indicate detected/annotated proteins; gray shading indicates all detected metabolic proteins. Data represent three independent biological replicates per age. Protein changes were assessed using moderated two-sided t tests with Benjamini–Hochberg FDR correction and classified as altered when q < 0.05 and |log₂ fold change| > 1. *q < 0.05, **q < 0.01, and ***q < 0.001.

**Figure S7| Mean lifespan of NAD⁺ pathway mutants and supplemented animals.**

Age at death of wild-type animals compared with *parp-1* (**S7A**), *parp-2* (**S7B**), *tank-1* (**S7C**), *tir-1* (**S7D**), and *pnc-1* (**S7E**) mutants, and of H₂O vehicle controls compared with animals receiving 1 mM NAM (**S7F**), 25 mM NAM (**S7G**), 1 mM NA (**S7H**), 25 mM NA (**S7I**), 1 mM trigonelline (**S7J**), or 25 mM trigonelline (**S7K**). (**S7L**) Age at death of H₂O-treated controls from two independent experiments. Percentages indicate the change in mean lifespan relative to the control. Boxplots show medians and IQRs, whiskers show the full range, and points represent individual animals. Data are from the cohorts in **Figure 7D and 7F**; group sizes are shown below each group. Statistical significance was determined by pairwise log-rank (Mantel–Cox) tests with Holm correction. **p < 0.01 and ***p < 0.001; unmarked comparisons were not significant.

**Supplemental Table 1:** *C. elegans* genes and encoded proteins (genes italicized lowercase; proteins uppercase, e.g., *act-1*/ACT-1)

| Gene | Protein | Encoded product |
|---|---|---|
| *afmd-1* | AFMD-1 | Arylformamidase (kynurenine formamidase) |
| *cdc-42* | CDC-42 | CDC42 Rho-family GTPase |
| *cyc-1* | CYC-1 | Cytochrome c1 |
| *haao-1* | HAAO-1 | 3-Hydroxyanthranilate 3,4-dioxygenase |
| *kmo-1* | KMO-1 | Kynurenine 3-monooxygenase |
| *kynu-1*(*flu-2*) | KYNU-1(FLU-2) | Kynureninase |
| *nkat-3* | NKAT-3 | Kynurenine aminotransferase |
| *nmat-1, nmat-2* | NMAT-1, NMAT-2 | Nicotinamide/nicotinic acid mononucleotide adenylyltransferase |
| *nmrk-1* | NMRK-1 | Nicotinamide riboside kinase |
| *nprt-1* | NPRT-1 | Nicotinate phosphoribosyltransferase |
| *parp-1, parp-2* | PARP-1, PARP-2 | Poly(ADP-ribose) polymerase 1 and 2 (formerly *pme-1*, *pme-2*) |
| *pmp-3* | PMP-3 | Peroxisomal membrane protein-related ABC transporter |
| *pnc-1, pnc-2* | PNC-1, PNC-2 | Nicotinamidase |
| *qns-1* | QNS-1 | Glutamine-dependent NAD<sup>+</sup> synthetase |
| *sir-2.1–sir-2.4* | SIR-2.1–SIR-2.4 | Sirtuins (Sir2-related NAD<sup>+</sup>-dependent deacylases) |
| *tank-1* | TANK-1 | Tankyrase |
| *tdo-2* | TDO-2 | Tryptophan 2,3-dioxygenase |
| *tir-1* | TIR-1 | TIR domain adaptor protein (SARM1 ortholog) |
| *umps-1* | UMPS-1 | Uridine monophosphate synthetase |

**Supplemental Table 2:** Strains used in this Study.

| *Organism* | Strain | Genotype | Source |
|---|---|---|---|
| *C. elegans* | N2 | WT | CGC |
| *C. elegans* | HV727 | pnc-1 (pk9605) IV | WHR |
| *C. elegans* | RB1042 | parp-1 (ok988) I | CGC |
| *C. elegans* | FX3401 | parp-2 (tm3401) II | WHR |
| *C. elegans* | RB684 | tank-1 (ok446) V | CGC |
| *C. elegans* | VC526 | tir-1 (gk264) III | CGC |
| *E. coli* | OP50 | uracil auxotroph, B strain of E. coli | CGC |

**Supplementary Table 3:**
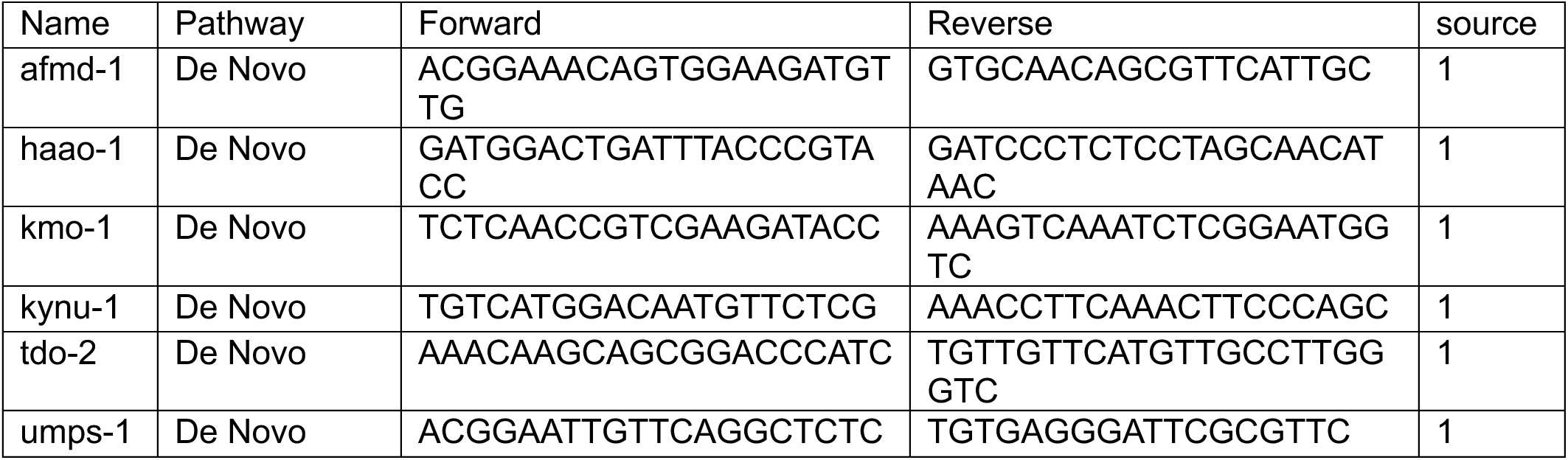
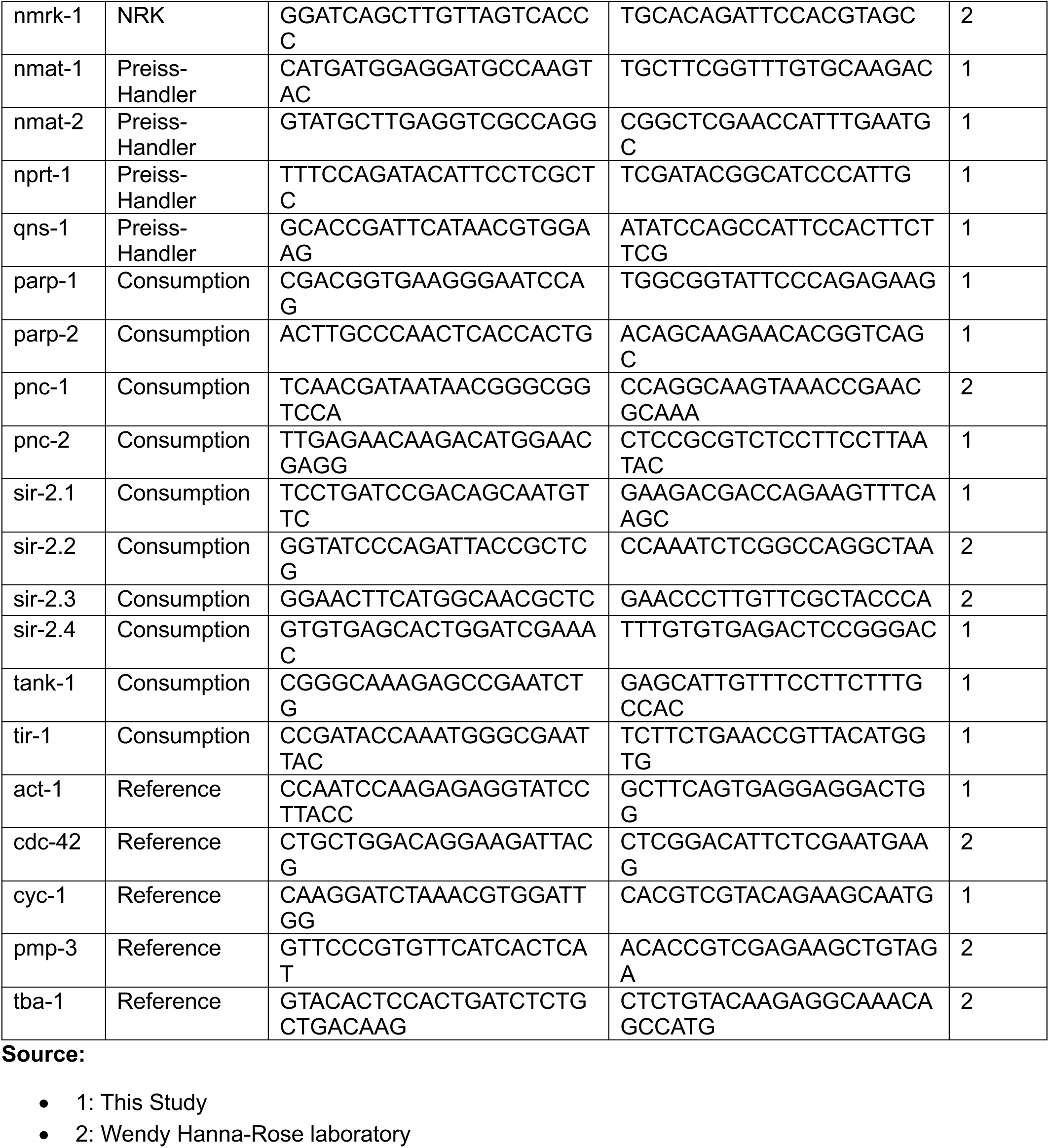
qRT-Primers.

## STAR METHODS

KEY RESOURCES TABLE.

| **REAGENT or RESOURCE** | **SOURCE** | **IDENTIFIER** |
|---|---|---|
| **Bacterial and virus strains** |  |  |
| E. coli OP50 | Caenorhabditis Genetics Center (CGC) | WormBase: OP50 |
| **Chemicals, peptides, and recombinant proteins** |  |  |
| Bacto Tryptone | Gibco (Thermo Fisher Scientific) | Cat# 211705 |
| Bacto Peptone | Thermo Fisher Scientific | Cat# 211677 |
| Yeast extract | Thermo Fisher Scientific | Cat# H26769.36 |
| Agar | Sigma-Aldrich | Cat# A7002 |
| Sodium chloride (NaCl) | Sigma-Aldrich | Cat# S9888 |
| Cholesterol (C27H46O) | Alfa Aesar | Cat# A11470-30 |
| Calcium chloride (CaCl2) | VWR | Cat# 1311-01 |
| Magnesium sulfate (MgSO4) | VWR | Cat# 2506-05 |
| Potassium phosphate monobasic (KH2PO4) | MilliporeSigma | Cat# PX1565 |
| Potassium phosphate dibasic (K2HPO4) | VWR | Cat# BDH9266-2.5KG |
| Sodium phosphate dibasic (Na2HPO4) | VWR | Cat# 97061-588 |
| Ammonium chloride (NH4Cl) | VWR | Cat# BDH9208-500G |
| Citric acid monohydrate | VWR | Cat# BDH9228 |
| Tri-potassium citrate monohydrate | Thermo Scientific Chemicals | Cat# AC244210010 |
| Ethylenediaminetetraacetic acid disodium salt dihydrate (Na2EDTA) | Invitrogen | Cat# 15576028 |
| Iron(II) sulfate heptahydrate (FeSO4·7H2O) | Thermo Scientific Chemicals | Cat# AC201392500 |
| Manganese(II) chloride tetrahydrate (MnCl2·4H2O) | VWR | Cat# 89230-036 |
| Zinc sulfate heptahydrate (ZnSO4·7H2O) | VWR | Cat# 470303-144 |
| Copper(II) sulfate pentahydrate (CuSO4·5H2O) | VWR | Cat# 470300-864 |
| TRIzol Reagent | Invitrogen | Cat# 15596026 |
| Nuclease-free water | VWR | Cat# 10220-402 |
| Chloroform | Sigma-Aldrich | Cat# C2432 |
| 2-Propanol (isopropanol) | Fisher Chemical | Cat# A426S-4 |
| Sodium acetate (CH3COONa), 3 M, pH 5.2 | EM Science | Cat# SX0265-1 |
| Phenol:chloroform:isoamyl alcohol (25:24:1), pH 8.0 | VWR | Cat# TS32711-1000 |
| Chloroform:isoamyl alcohol (24:1) | VWR | Cat# X205-450ML |
| GlycoBlue Coprecipitant | Invitrogen | Cat# AM9515 |
| Water, HPLC grade | VWR Chemicals BDH | Cat# BDH23595.400 |
| 2-Propanol, HPLC grade | VWR Chemicals BDH | Cat# BDH20880.400 |
| Acetonitrile, Optima LC/MS grade | Fisher Chemical | Cat# A955-4 |
| Methanol, HPLC grade | Fisher Chemical | Cat# A452-4 |
| Formic acid, Optima LC/MS grade | Fisher Chemical | Cat# A117-50 |
| Ammonium bicarbonate | MilliporeSigma | Cat# A6141 |
| Ammonium acetate, LC-MS Ultra | Honeywell Fluka | Cat# 14267 |
| Ammonium hydroxide, 35% for HPLC | Thermo Scientific Chemicals | Cat# AC460801000 |
| Nicotinic acid | Sigma-Aldrich | Cat# 72309 |
| Nicotinamide | Sigma-Aldrich | Cat# 72340 |
| Trigonelline hydrochloride | Sigma-Aldrich | Cat# T5509 |
| Nicotinic acid-2,4,5,6-d4 | Cambridge Isotope Laboratories | Cat# DLM-4578-1 |
| Nicotinamide-2,4,5,6-d4 | Cambridge Isotope Laboratories | Cat# DLM-6883 |
| **Critical commercial assays** |  |  |
| EasyPep MS Sample Prep Kit | Thermo Fisher Scientific | Cat# A45733 |
| Pierce Quantitative Colorimetric Peptide Assay | Thermo Fisher Scientific | Cat# 23275 |
| Pierce BCA Protein Assay Kit | Thermo Fisher Scientific | Cat# 23227 |
| qScript cDNA Synthesis Kit | Quantabio | Cat# 95047 |
| **Deposited data** |  |  |
| Metabolomics and stable-isotope tracing data | This paper | *Available Upon Request* |
| Proteomics data | This paper | *Available Upon Request* |
| Original code (R scripts) | This paper | *Available Upon Request* |
| **Experimental models: Organisms/strains** |  |  |
| C. elegans: Strain N2 (Bristol) | Caenorhabditis Genetics Center (CGC) | WormBase: N2 |
| C. elegans: Strain HV727: pnc-1(pk9605) IV | W. Hanna-Rose laboratory, Penn State | WormBase: HV727 |
| C. elegans: Strain RB1042: parp-1(ok988) I | Caenorhabditis Genetics Center (CGC) | WormBase: RB1042 |
| C. elegans: Strain FX3401: parp-2(tm3401) II | W. Hanna-Rose laboratory, Penn State | WormBase: FX3401 |
| C. elegans: Strain RB684: tank-1(ok446) V | Caenorhabditis Genetics Center (CGC) | WormBase: RB684 |
| C. elegans: Strain VC526: tir-1(gk264) III | Caenorhabditis Genetics Center (CGC) | WormBase: VC526 |
| **Oligonucleotides** |  |  |
| Primers for qRT-PCR, see Table S1 | This paper; W. Hanna-Rose laboratory | N/A |
| **Software and algorithms** |  |  |
| WormLab 2021-2026 | MBF Bioscience | RRID:SCR_017669; https://www.mbfbioscience.com/wormlab |
| El-MAVEN v0.12.0 | Melamud et al., 2010; Agrawal et al., 2019 | RRID:SCR_022159; https://elucidata.io/el-maven/ |
| AccuCor | Su et al., 2017 | https://cran.r-project.org/package=accucor |
| MSConvert, ProteoWizard v3.0.25100 | Chambers et al., 2012 | RRID:SCR_012056; https://proteowizard.sourceforge.io/ |
| DEP2 v0.5.28.2 | Feng et al., 2023 | https://github.com/mildpiggy/DEP2 |
| MaxQuant v2.8.1.0 | Cox and Mann, 2008 | RRID:SCR_014485; https://www.maxquant.org/ |
| GraphPad Prism 10.4.0 | GraphPad Software | RRID:SCR_002798; https://www.graphpad.com |
| R v4.4.3 | R Core Team | RRID:SCR_001905; https://www.r-project.org/ |
| RStudio v2026.08.2+200 | Posit Software, PBC | RRID:SCR_000432; https://posit.co/products/open-source/rstudio/ |
| **Other** |  |  |
| C. elegans Synchronizer | NemaSync | Model CES-700 |
| Bead mill tubes, 2 mL | VWR | Cat# 10158-556 |
| Zirconium oxide ceramic beads, 2.8 mm | Fisherbrand | Cat# 15-340-160 |
| Zirconia/silica beads, 1.0 mm | Fisher Scientific | Cat# NC9847287 |
| XBridge BEH Amide column (2.1 × 150 mm, 3.5 µm) | Waters | Cat# 186004861 |
| Orbitrap Exploris 240 mass spectrometer | Thermo Fisher Scientific | N/A |
| Orbitrap Eclipse Tribrid mass spectrometer | Thermo Fisher Scientific | N/A |
| Vanquish Neo UHPLC system | Thermo Fisher Scientific | N/A |
| PepMap Neo C18 column (75 µm × 150 mm, 2 µm) | Thermo Fisher Scientific | Cat# DNV75150PN |
| PepMap Neo C18 trap cartridge (0.3 × 5 mm, 5 µm) | Thermo Fisher Scientific | Cat# 174500 |
| Varioskan LUX multimode microplate reader | Thermo Fisher Scientific | N/A |
| Basler acA2440 camera | Basler | Model acA2440-35uM |

### *C. elegans* culture and strains

Wild-type N2 (Bristol) animals and other strains were obtained from the *Caenorhabditis* Genetics Center (CGC, University of Minnesota) and Wendy Hanna-Rose lab at Penn State (Supplemental Table 2). Worms were maintained at 20 °C on nematode growth medium (NGM) plates seeded with *Escherichia coli* OP50 using standard procedures **[84]**. Animals were maintained under well-fed conditions, and bacterial lawns were replenished before depletion to avoid starvation-associated changes in metabolic state.

Synchronized cohorts were generated by passing mixed-stage populations through a NemaSync microfiltration system, collecting L1 larvae, and transferring them to freshly seeded NGM plates. Aging cohorts were maintained free of progeny by daily washing and gravity-based separation of adults from smaller progeny, followed by transfer of the adults to freshly seeded plates. This physical separation procedure avoided the use of reproductive inhibitors, including 5-fluoro-2′-deoxyuridine (FUdR), and sterilizing mutations. Animals were collected at 62 h (D03) and on Days 8 (D08), 12 (D12), and 15 (D15).

### Lifespan analysis

Lifespan assays were performed at 20 °C on NGM plates seeded with 400 µL of *E. coli* OP50. Each assay was repeated in at least three independent experiments. Cohorts were synchronized by timed egg laying, and 25 L4 animals were placed on each assay plate at the start of the experiment, designated day 0. Animals were transferred to fresh plates daily.

Survival was assessed daily by observing spontaneous movement or, for older animals, by gently touching them with a platinum wire. Animals that did not respond were scored as dead. Animals that underwent internal hatching (“bagging”) or vulval rupture, left the plate, or desiccated on the plate wall were censored at the time of observation.

### Brood assay

Progeny produced by animals used in the lifespan assays were retained on the plates after parental transfer and maintained at 20 °C. Two days after removal of the parent, each plate was imaged using an iPad mounted to a microscope. Progeny was scored at the L3-to-young-adult stages, when animals were sufficiently large for reliable counting and had not yet begun reproducing. Total brood size was calculated as the sum of viable progeny produced by each individual over its reproductive period.

### Crawling-speed analysis (WormLab)

Crawling speed was measured using WormLab software (MBF Bioscience) at the indicated ages. Animals were recorded for 1 min on clean, unseeded NGM plates at room temperature. Videos were acquired against a light background using a Nikon lens mounted on a Basler acA2440 camera at 2456 × 2052 pixels in Mono8 mode and 14 frames per second.

For each recording, an intensity threshold was set to distinguish animals from the background. WormLab then identified and tracked individual animals using specified body-length and body-width ranges **[85]**. Crawling speed was defined as distance traveled per second along the animal’s central axis and calculated frame by frame. To reduce tracking artifacts, frame-wise speeds exceeding 500 µm/s, which typically arise from temporary loss and reacquisition of the animal, were excluded, and the remaining values were averaged for each animal **[85]**.

### Thrashing assay

Animals were maintained at 20 °C from the L4 stage and assayed at the indicated ages. Individual worms were transferred to a 15 µL drop of M9 buffer on a clean, unseeded NGM plate at room temperature. Following 1 min of acclimation, swimming worms were recorded for 1 min using a Basler acA2440 camera and the WormLab system (MBF Bioscience). One thrash was defined as a complete change in the direction of bending at the mid-body. Each animal was scored once; the number of thrashes was counted manually from the recorded videos. Assays included 20–30 animals per condition.

### Sample collection and metabolite extraction

Worms were washed three times with autoclaved Milli-Q water to remove adherent bacteria, pelleted, snap-frozen in liquid nitrogen, and stored at −80 °C until extraction. Frozen pellets were thawed on wet ice, and 25 µL aliquots of worm slurry were transferred to prechilled 2.0 mL bead-mill tubes containing 6–10 homogenization beads and 1.0 mL of ice-cold extraction solvent. The solvent consisted of ACN:MeOH:H_2_O (40:40:20, v/v/v; LC-MS/Optima grade) supplemented with 0.5% (v/v) formic acid. Samples were processed in triplicate. An extraction blank containing beads and solvent without biological material was processed in parallel.

Samples were homogenized at 4 °C using two 30 s bead-milling cycles at 6.0 m/s, separated by a 15 s interval. Homogenates were immediately neutralized with 88 µL of 15.8% (w/v) ammonium bicarbonate, briefly vortexed, and incubated at −20 °C for 15 min to promote protein precipitation. Samples were centrifuged for 20 min at 4 °C, and 800 µL of clarified supernatant was transferred to clean microcentrifuge tubes. A 100 µL aliquot was reserved for protein quantification using a bicinchoninic acid (BCA) assay.

### Sample preparation for HILIC–LC-MS

For hydrophilic interaction liquid chromatography–mass spectrometry (HILIC–LC-MS), 200 µL of extract was evaporated to dryness under a gentle stream of nitrogen gas. Residues were reconstituted in 50 µL of ACN:MeOH:H_2_O (40:40:20, v/v/v) by gentle pipetting and centrifuged for 20 min at 4 °C. A 40 µL aliquot of clarified extract was transferred to a glass autosampler vial for analysis in positive and negative electrospray ionization modes. Remaining extracts were stored at −80 °C.

### Stable-isotope tracing

Stable-isotope labeling was performed immediately before sample collection at D03, D08, D12, and D15. Animals were recovered from NGM plates in 1 mL of M9 buffer and pelleted by centrifugation at 500 × *g* for 1 min. Each pellet was resuspended in 1 mL of concentrated, heat-killed *E. coli* OP50. D₄-nicotinic acid (D₄-NA) or D₄-nicotinamide (D₄-NAM) was added as 100 µL of a 200 mM stock, and M9 buffer was added to a final volume of 2 mL, yielding a tracer concentration of 10 mM. OP50 was autoclaved before use to prevent bacterial metabolism of the tracers during incubation. Cultures were incubated at room temperature for 4 h with gentle rocking. Pilot experiments using shorter and longer incubations were conducted to assess the linearity of label incorporation over this interval.

Following incubation, animals were pelleted at 500 × *g* for 1 min. A 1.5 mL aliquot of the extracellular medium was transferred to a separate tube and retained for LC-MS analysis. Worm pellets were washed three times with 15 mL of autoclaved Milli-Q water, with centrifugation between washes, then snap-frozen in liquid nitrogen and stored at −80 °C until extraction. Water was used for the final washes to minimize ion suppression from residual M9 salts. Isotopologue distributions were corrected for natural isotope abundance using AccuCor **[86]**.

### Metabolite measurement

Extracts were analyzed within 24 h using hydrophilic interaction liquid chromatography coupled to an Orbitrap Exploris 240 mass spectrometer (Thermo Scientific), as previously described **[87]**. Metabolites were separated on an XBridge BEH Amide column (2.1 × 150 mm, 3.5 µm particle size; Waters, Milford, MA). Mobile phase A consisted of H_2_O:ACN (95:5, v/v), and mobile phase B consisted of H_2_O:ACN (10:90, v/v), each containing 20 mM ammonium acetate and 20 mM ammonium hydroxide.

The gradient was programmed as follows: 90% B at 0–2 min; 75% B at 3–7 min; 70% B at 8–9 min; 50% B at 10–12 min; 25% B at 13–14 min; 0% B at 16–20 min; and 90% B at 21–25 min. The flow rate was 150 µL/min, the column temperature was 25 °C, and the autosampler temperature was 5 °C. The injection volume was 5 µL.

Mass spectra were acquired in positive and negative ionization modes at a resolving power of 180,000 at *m/z* 200, with an automatic gain control (AGC) target of 3 × 10⁶, a maximum injection time of 30 ms, and a scan range of *m/z* 70–1000. Raw files were converted to mzML format using the msconvert command-line utility **[88]** and processed in EL-MAVEN version 12 **[89]**.

### RNA isolation and quantitative reverse-transcription PCR

Approximately 200–250 synchronized animals were collected per biological sample at each age, washed three times with M9 buffer to remove bacteria, and resuspended in 250 µL TRIzol before snap-freezing in liquid nitrogen. Worms were homogenized in bead mill tubes containing ceramic beads, and total RNA was extracted using TRIzol reagent (Thermo Fisher Scientific, Carlsbad, CA) according to the manufacturer’s instructions, followed by chloroform extraction and centrifugation at 12,700 rpm for 15 min at 4°C. The aqueous phase was recovered, RNA was precipitated with 2-propanol and GlycoBlue, pelleted, washed with 70% ethanol, air-dried, and resuspended in RNase-free water. All samples were treated with DNase I-XT to remove genomic DNA and further purified by phenol:chloroform extraction and ethanol precipitation. RNA concentration and purity were assessed using a Varioskan LUX plate reader (Thermo Fisher Scientific). For each sample, 1.0 µg of RNA was reverse transcribed using the qScript cDNA Synthesis Kit (Quanta Biosciences, Beverly, MA) by following the manufacturer’s instructions.

Quantitative PCR was performed in 20 µL reactions containing cDNA, gene-specific primers, and PerfeCTa [confirm exact master mix] (Quanta Biosciences). Cycling conditions consisted of 95 °C for 3 min, followed by 40 cycles of 95 °C for 15 s and 60 °C for 1 min. Each biological sample was analyzed in technical triplicate, with six independent biological replicates per age. No-template and no-reverse-transcriptase controls were included on every plate.

Relative expression was calculated using the 2^−ΔΔCt method **[90]**, with D03 as the reference age and normalization to the geometric mean of three reference genes (*cdc-42, cyc-1, and pmp-3*) **[91]**. Expression values were reported as fold changes or log₂ fold changes relative to D03, as indicated. Primer sequences are provided in Supplementary Table 3.

### Proteomic sample preparation and peptide purification

Wild-type N2 populations were synchronized by a 30 min timed egg lay. Three independent biological replicates were collected per age, with 50 animals pooled per D03 sample and 20 animals pooled per D08, D12, or D15 sample. Animals were collected directly into kit lysis buffer and immediately frozen at −80 °C.

Protein extraction, digestion, and peptide purification were performed using the EasyPep™ MS Sample Prep Kit (Thermo Fisher Scientific; catalog no. A45733) according to the manufacturer’s instructions. Peptide concentrations were measured using the Pierce™ Quantitative Peptide Assay (Thermo Fisher Scientific; catalog no. 23295), with absorbance recorded at 562 nm on a Varioskan LUX plate reader. A total of 500 ng of purified peptides were submitted to the Proteomics and Mass Spectrometry Core Facility at The Pennsylvania State University for LC-MS/MS analysis.

### Proteomics (Nano-Liquid Chromatography – Mass Spectrometry)

The dried down peptides were dissolved in 10 mL (please check if this is correct volume) of 2% acetonitrile, 0.1% formic acid, and 5 mL of this solution were used for analysis.

Peptides were separated on a Thermo DNV PepMap Neo reverse-phase column (150 mm x 75 μm, 2 μm, 100 A) with a 5 mm x 0.3 mm PepMap Neo C18 5 μm trapping cartridge (Thermo). The following gradient was delivered by a Vanquish Neo nano-LC system (Thermo): 0-80 min 1% – 40% B, 80-100 min 40% – 70% B, a 10-min column wash at 95% B, followed by the column equilibration at 1% B. Mobile phase A was 0.1% formic acid in water, mobile phase B was 80% aqueous acetonitrile, containing 0.1% formic acid; the flow rate was 300 nL/min.

The mass spectra were acquired on a Thermo Orbitrap Eclipse instrument using a method based on the Universal template. The MS scans were acquired at R 120,000 over 375 – 1500 m/z range; automatic gain control (AGC) target was set to 4 × e5 and the maximum inject time was set to Auto (max. 246 ms). The data-dependent precursor selection for MS2 had a cycle time of 3 s. The MS2 precursors were isolated in the quadrupole (0.7 m/z window) and analyzed in the ion trap. CID collision energy was 35%, AGC target 2 × e4, maximum inject time was 35 ms. Scan range was set from 150 to 1500 m/z.

### Proteomics data analysis

Raw files were processed in MaxQuant (v2.8.1.0) **[92]** using the integrated Andromeda search engine **[93]**. Spectra were searched against three FASTA files: the *C. elegans* reference proteome from UniProt (UP000001940, taxon 6239; downloaded July 2026), the *C. elegans* protein set from WormBase ParaSite release 19 (BioProject PRJNA13758, WBPS19) **[94]**, and the universal protein contaminant library **[95]** (release 0602). The MaxQuant built-in contaminant list was also included. Because the UniProt and WormBase sequence sets overlap, proteins that shared identical identified peptides were collapsed into single protein groups by MaxQuant. Searches specified trypsin/P with up to two missed cleavages, carbamidomethylation of cysteine as a fixed modification, and oxidation of methionine and protein N-terminal acetylation as variable modifications. Because MS2 spectra were acquired in the ion trap, the MS/MS match tolerance was set to 0.5 Da. Peptide-spectrum match and protein false discovery rates were set to 1% using a target-decoy approach. Proteins were quantified using the MaxLFQ algorithm **[96]**, with match between runs enabled.

The MaxQuant proteinGroups output was analyzed using the web-based DEP2 Shiny application (accessed July 2026) **[97]**. LFQ intensity columns were used for quantification, with gene names as protein labels and protein IDs as identifiers. Protein groups flagged as potential contaminants were removed, as were reverse-database hits and proteins identified only by site. Proteins with no missing values in all three biological replicates of at least one age were retained, yielding 1,810 quantified proteins. LFQ intensities were log₂-transformed and quantile-normalized **[98]**, and remaining missing values were imputed by Bayesian principal component analysis (BPCA) **[99]**. Differential abundance at D08, D12, and D15 relative to D03 was assessed using limma-based moderated two-sided t tests **[100]** with Benjamini–Hochberg correction. Proteins were considered significantly altered when they met both criteria, q < 0.05 and |log₂ fold change| > 1.

### Supplementation

Filter-sterilized stock solutions of nicotinamide (Sigma, catalog no. 72340), nicotinic acid (Sigma, catalog no. 72309), and trigonelline hydrochloride (Sigma, catalog no. T5509) were added to plates seeded with UV-killed *E. coli* OP50 to reach final concentrations of 1 mM and 25 mM depending on the experiment. The supplemented plates were incubated at room temperature for 24 hours prior to use. Wild-type N2 populations were synchronized via a 1-hour timed egg lay directly onto the supplemented plates, where the resulting larvae were allowed to hatch and develop. Following development, L4/young adult hermaphrodites were singled and assayed for lifespan and progeny production for the indicated phenotypes.

### Statistical analysis

Analyses were performed using GraphPad Prism version 10.4.0 (GraphPad Software) and additional software specified for individual analyses. Sample sizes, measures of central tendency and variability, and statistical tests are reported in the corresponding figure legends. Biological replicates were treated as independent samples; technical injections were averaged within each biological sample before statistical analysis. For locomotion and brood assays, individual animals were the observational units.

Age-associated differences in transcript expression, metabolite abundance, isotope enrichment, and derived metabolite ratios were evaluated by one-way ANOVA, with transformations specified in the figure legends. Comparisons with D03 used Dunnett’s multiple-comparison test for transcript expression and Tukey’s honestly significant difference test for metabolomic and tracing analyses. Locomotion and brood-size outcomes were analyzed by one-way ANOVA followed by Dunnett’s test against wild-type animals or the vehicle control. Overall age effects for proteomic modules were assessed by one-way ANOVA. Selected metabolite ratios in **Figure 6F** were compared with D03 using unpaired, two-tailed Welch’s *t* tests.

Differential protein abundance was assessed using moderated two-sided *t* tests, with Benjamini–Hochberg false discovery rate (FDR) correction across all 1,810 quantified proteins. FDR correction was also applied to the metabolite and metabolite-ratio analyses indicated in the figure legends. Adjusted values are reported as *q* values; unadjusted values are reported as *p* values. Fold-change thresholds, where applied, are specified for each analysis. Transcript estimates in **Figure 1D** were obtained using a Poisson-lognormal generalized linear mixed model and are presented as posterior means with 95% highest posterior density credible intervals. An effect was considered credible when its interval excluded zero.

Survival distributions were compared using overall log-rank (Mantel–Cox) tests followed by pairwise log-rank tests with Holm correction for multiple comparisons. Median lifespans and the numbers of animals scored and censored are reported for each condition. Observations flagged by Rosner’s generalized extreme Studentized deviate test in **Figure 7** were retained in the analyses.

Statistical significance was defined as *p* < 0.05 or FDR-adjusted *q* < 0.05, as applicable. Asterisks indicate significance levels defined in each figure legend; ns denotes a nonsignificant comparison. Sample sizes refer to independent biological samples or individual animals, as specified, rather than technical measurements. Descriptive pathway summaries calculated across metabolites were not used for inferential testing.

