## Supplemental Figures for "Coordinated remodeling of NAD⁺ metabolic networks across the lifespan of *Caenorhabditis elegans*"

### Supplementary Figure 1

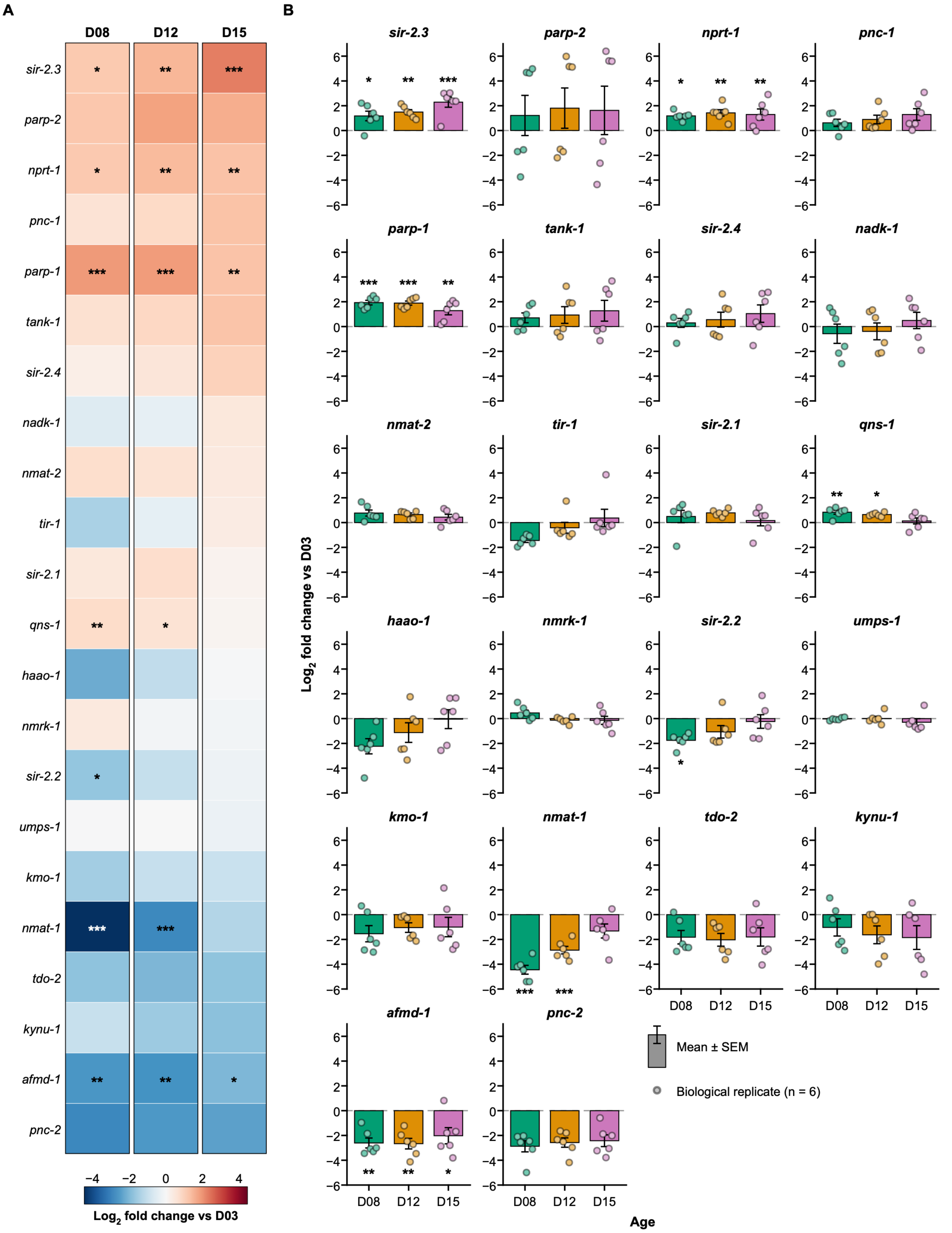

**Figure S1| Pathway-level and replicate-level analysis of $NAD^+$-related transcripts.**

(**S1A**) Heatmap of mean $\log_2$ fold changes relative to D03 for the 22 $NAD^+$-related genes in Figure 1C at D08, D12, and D15, grouped by pathway: de novo, Preiss–Handler, NR kinase (NRK), salvage, sirtuins, PARPs/TIR, and $NADP^+$ synthesis. (**S1B**) $\text{Log}_2$ fold changes relative to D03 for each gene. Bars show mean ± SEM; points represent individual biological replicates. Expression was measured by qRT-PCR and normalized to the geometric mean of *cdc-42*, *cyc-1*, and *pmp-3*. Data represents six independent biological replicates per age. Statistical significance was determined by one-way ANOVA followed by Dunnett's multiple-comparison test against D03. *$p < 0.05$, **$p < 0.01$, and ***$p < 0.001$.

### Supplementary Figure 2

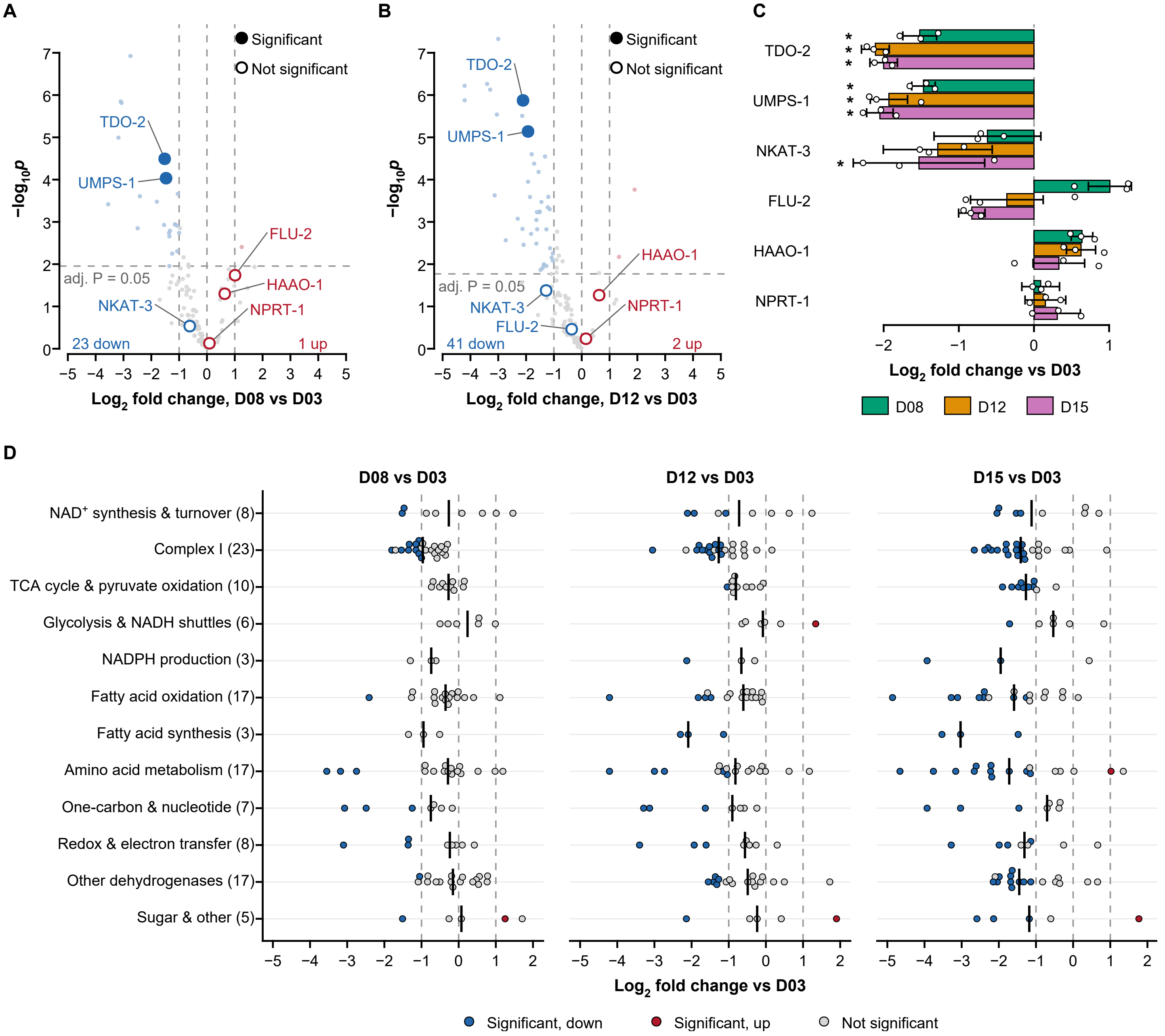

**Figure S2| Progressive decline of the NAD$^+$-associated proteome across adulthood.**

(**S2A and S2B**) Volcano plots of protein-abundance changes at D08 (**S2A**) and D12 (**S2B**) relative to D03 for 124 NAD$^+$-related and NAD$^+$-dependent proteins. Vertical dashed lines indicate $|\log_2$ fold change$|$ = 1. Twenty-three and 41 proteins decreased significantly at D08 and D12, respectively, and one and two proteins increased. Labeled NAD$^+$ biosynthetic enzymes are shown as filled (both criteria met) or open (criteria not met) circles. (**S2C**) $\text{Log}_2$ fold changes relative to D03 for TDO-2, UMPS-1, NKAT-3, FLU-2, HAAO-1, and NPRT-1. Bars show mean ± SD; points represent individual biological replicates. (**S2D**) $\text{Log}_2$ fold changes relative to D03 for the 124 proteins grouped into 12 functional categories. Numbers in parentheses indicate proteins per category, and vertical black lines indicate category medians. Blue and red points indicate significantly decreased and increased proteins, respectively; gray points indicate proteins that did not meet both criteria. Data represent three independent biological replicates per age. Statistical significance was assessed using moderated two-sided t tests with Benjamini–Hochberg FDR correction across all 1,810 quantified proteins. Proteins were classified as altered when $q < 0.05$ and $|\log_2$ fold change$| > 1$; asterisks in (**S2C**) indicate that both criteria were met.

Supplementary Figure 3

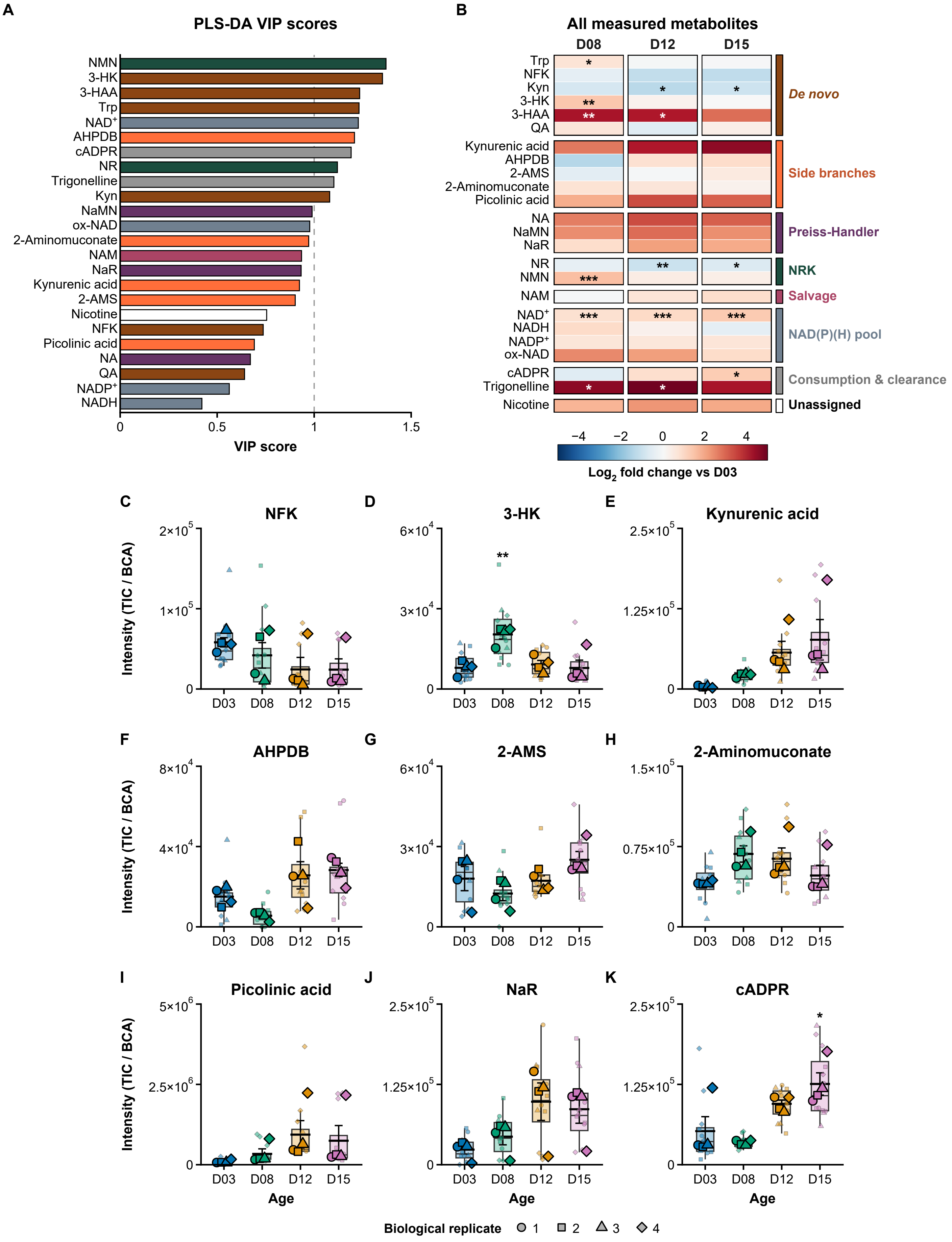

**Figure S3| Extended profiling of the steady-state NAD$^+$ metabolome.**

(**S3A**) Variable importance in projection (VIP) scores from the PLS-DA model in Figure 3A. Bar colors indicate pathway assignment as in (B); the dashed line marks VIP = 1. (**S3B**) Heatmap of mean $\log_2$ fold changes relative to D03 for all 24 measured metabolites at D08, D12, and D15, grouped by pathway assignment. (**S3C–K**) Normalized abundance of N-formylkynurenine (NFK; **S3C**), 3-hydroxykynurenine (3-HK; **S3D**), kynurenic acid (**S3E**), 4-(2-Amino-3-hydroxyphenyl)-2,4-dioxobutanoate (AHPDB; **S3F**), 2-aminomuconate semialdehyde (2-AMS; **S3G**), 2-aminomuconate (**S3H**), picolinic acid (**S3I**), nicotinic acid riboside (NaR; **S3J**), and cyclic ADP-ribose (cADPR; **S3K**). Metabolite signals were normalized to total ion count and BCA-measured protein abundance. Boxplots show medians and IQRs, with whiskers extending to 1.5 × IQR. Large, outlined symbols represent the mean of technical injections for each biological replicate; smaller translucent symbols represent individual injections. Data represent four independent biological replicates per age. Statistical significance was determined by one-way ANOVA followed by Tukey's HSD test; asterisks in (**S3B–K**) indicate significant differences from D03. *$p < 0.05$, **$p < 0.01$, and ***$p < 0.001$.

### Supplementary Figure 4

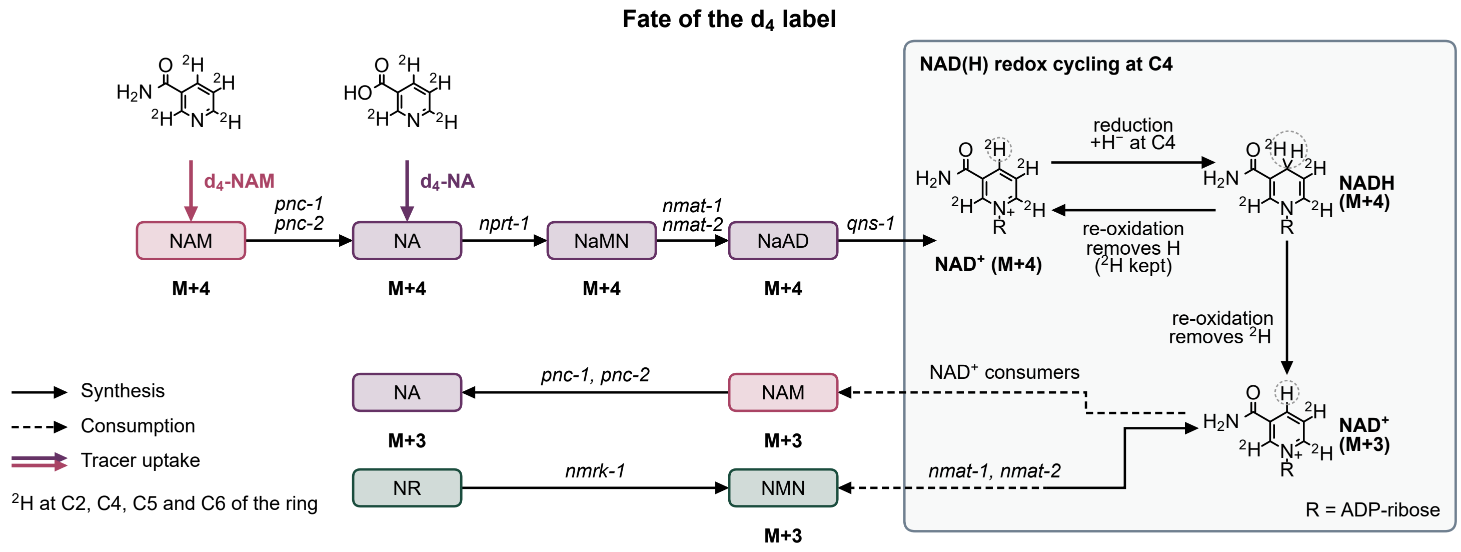

**Figure S4| Fate of the deuterium label from $D_4$-NAM and $D_4$-NA in the $NAD^+$ network.**

(**S4A**) $D_4$-NAM and $D_4$-NA carry deuterium ($^2H$) at C2, C4, C5, and C6 of the pyridine ring. $D_4$-NAM is deamidated to $D_4$-NA by PNC-1 and PNC-2, and both tracers enter $NAD^+$ through the Preiss–Handler pathway (NPRT-1, NMAT-1/NMAT-2, and QNS-1), generating M+4 $NAD^+$. Reduction of $NAD^+$ to NADH adds an unlabeled hydride at C4. Re-oxidation removes either the unlabeled hydrogen, regenerating M+4 $NAD^+$, or the C4 deuterium, generating M+3 $NAD^+$. $NAD^+$ consumers release M+3 NAM, which can be deamidated to M+3 NA. M+3 NMN arises from $NAD^+$ turnover and is converted to $NAD^+$ by NMAT-1/NMAT-2; NR enters through NMRK-1. Solid arrows indicate synthesis, dashed arrows indicate consumption, and double arrows indicate tracer uptake. R, ADP-ribose.

**Supplementary Figure 5**

Biological replicate ● 1 ■ 2 ▲ 3

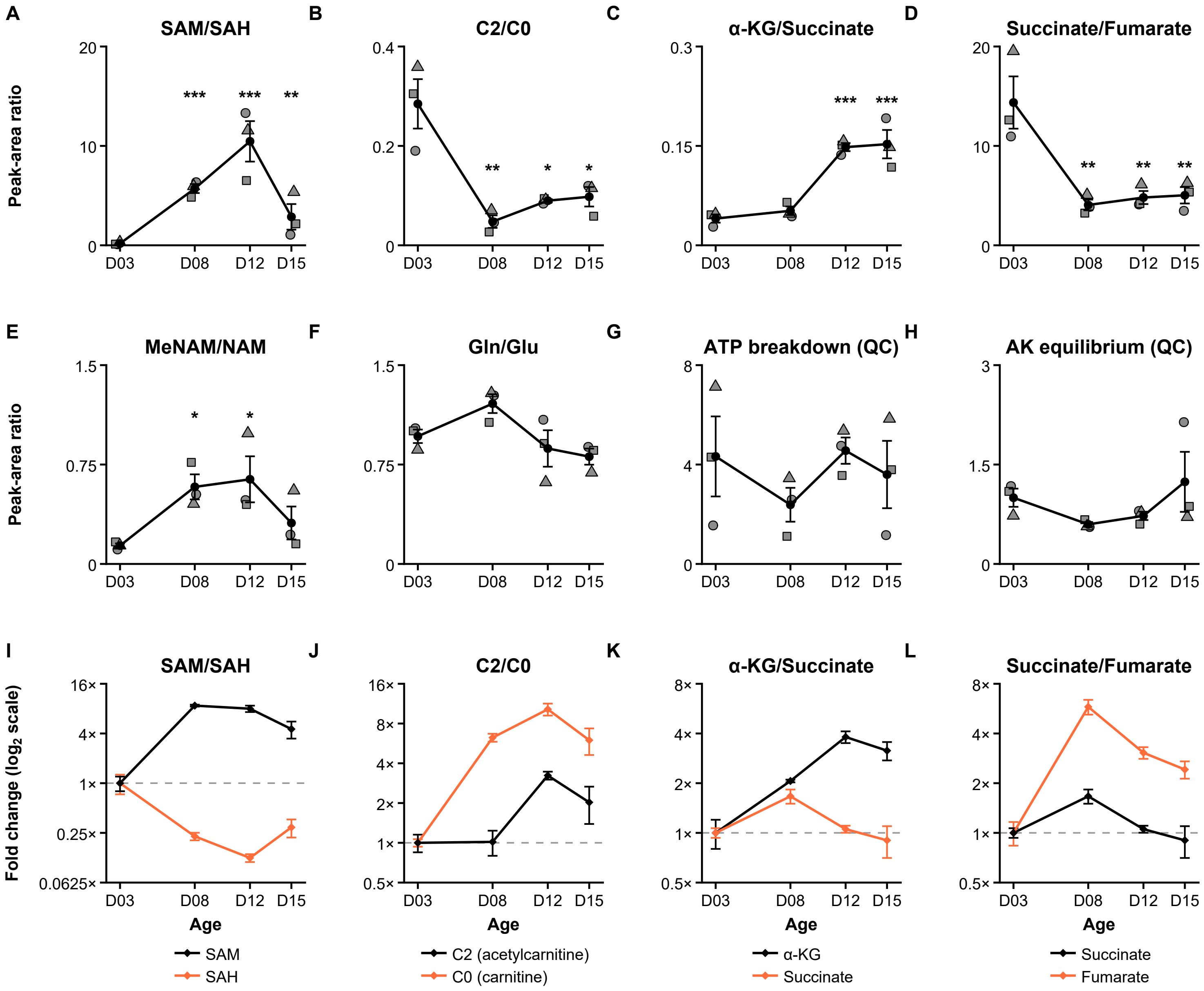

**Figure S5| Additional metabolite ratios reporting on methylation, carnitine, and TCA cycle status.**

(**S5A–F**) Peak-area ratios of S-adenosylmethionine/S-adenosylhomocysteine (SAM/SAH; **S5A**), acetylcarnitine/free carnitine (C2/C0; **S5B**), α-ketoglutarate/succinate (**S5C**), succinate/fumarate (**S5D**), $N^1$-methylnicotinamide/nicotinamide (MeNAM/NAM; **S5E**), and glutamine/glutamate (**S5F**). (**S5G and S5H**) Quality-control ratios for ATP breakdown (**S5G**) and adenylate kinase (AK) equilibrium (**S5H**). (**S5I– S5L**) Fold changes of the individual metabolites contributing to the ratios in (**S5A– S5D**), normalized to the D03 mean and displayed on a $\log_2$ scale. Black symbols and lines show mean ± SEM; gray symbols in (**S5A–S5H**) represent individual biological replicates. Data represent three independent biological replicates per age. Values are semi-quantitative LC-MS peak-area ratios from whole-animal extracts. Statistical significance was determined by one-way ANOVA of $\log_2$-transformed ratios followed by Tukey's HSD test; asterisks indicate significant differences from D03. *$p < 0.05$, **$p < 0.01$, and ***$p < 0.001$.

### Supplementary Figure 6

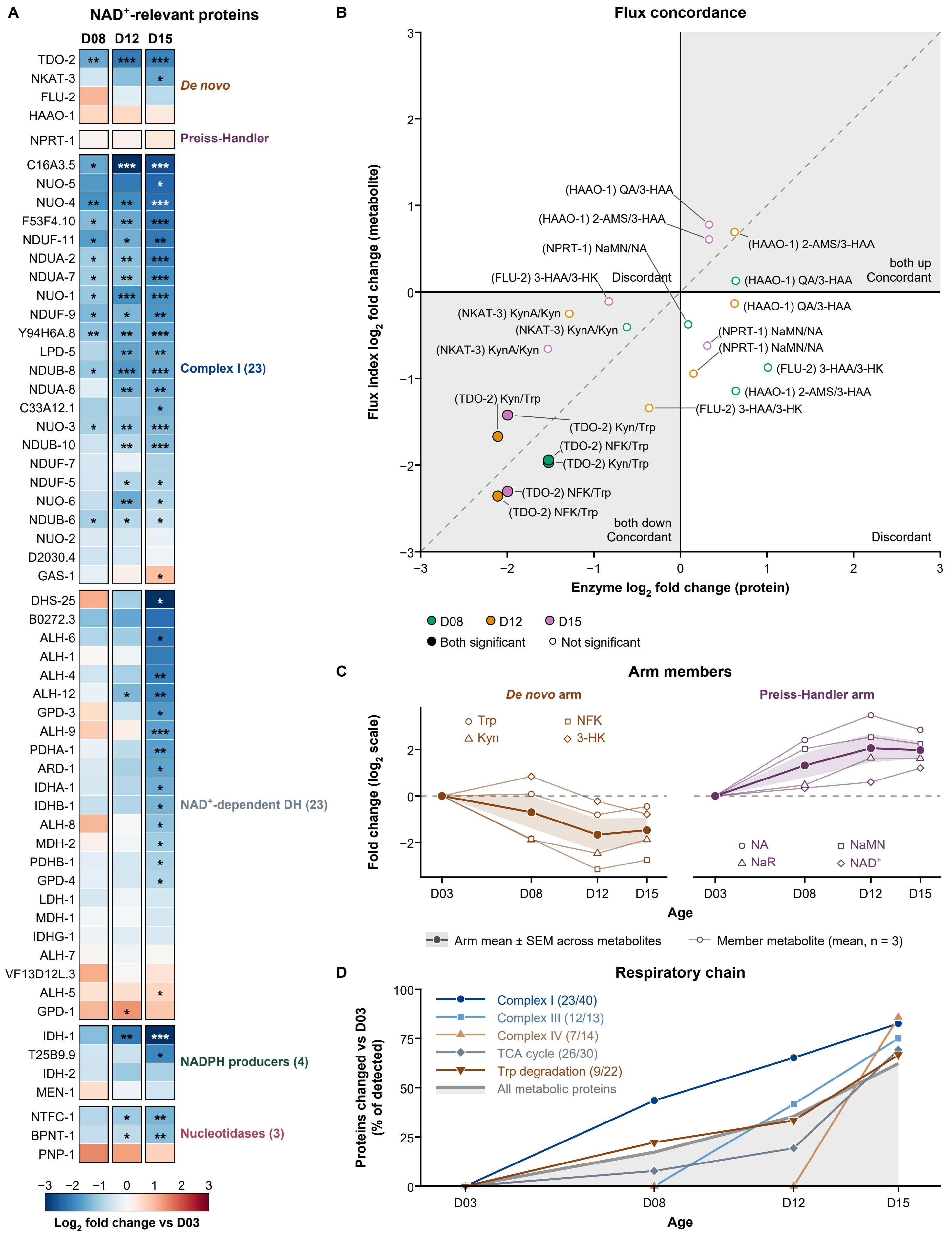

**Figure S6| Protein-level and pathway-level details of the multi-omic integration.**

(**S6A**) Heatmap of $\log_2$ fold changes relative to D03 for $NAD^+$-relevant proteins at D08, D12, and D15, grouped into de novo, Preiss–Handler, Complex I (23 proteins), $NAD^+$-dependent dehydrogenase (23), NADPH-producing (4), and nucleotidase (3) modules. (**S6B**) Labeled version of Figure 6E comparing enzyme $\log_2$ fold changes with changes in the corresponding product-to-substrate metabolite ratios. Each point is labeled with its enzyme and ratio; colors indicate age. Filled circles indicate that both changes were significant; open circles indicate that at least one was not. Shaded quadrants indicate concordant changes, and the dashed diagonal indicates the 1:1 line. (**S6C**) Trajectories of individual metabolites in the de novo arm (Trp, NFK, Kyn, and 3-HK) and Preiss–Handler arm (NA, NaMN, NaR, and $NAD^+$) summarized in Figure 6A. Open symbols show member means; filled symbols and shading show the arm mean ± SEM across metabolites. (**S6D**) Percentage of detected proteins differing significantly from D03 in Complex I, Complex III, Complex IV, the TCA cycle, and tryptophan degradation. Numbers in parentheses indicate detected/annotated proteins; gray shading indicates all detected metabolic proteins. Data represent three independent biological replicates per age. Protein changes were assessed using moderated two-sided t tests with Benjamini–Hochberg FDR correction and classified as altered when $q < 0.05$ and $|\log_2 \text{fold change}| > 1$. *$q < 0.05$, **$q < 0.01$, and ***$q < 0.001$.

Supplementary Figure 7

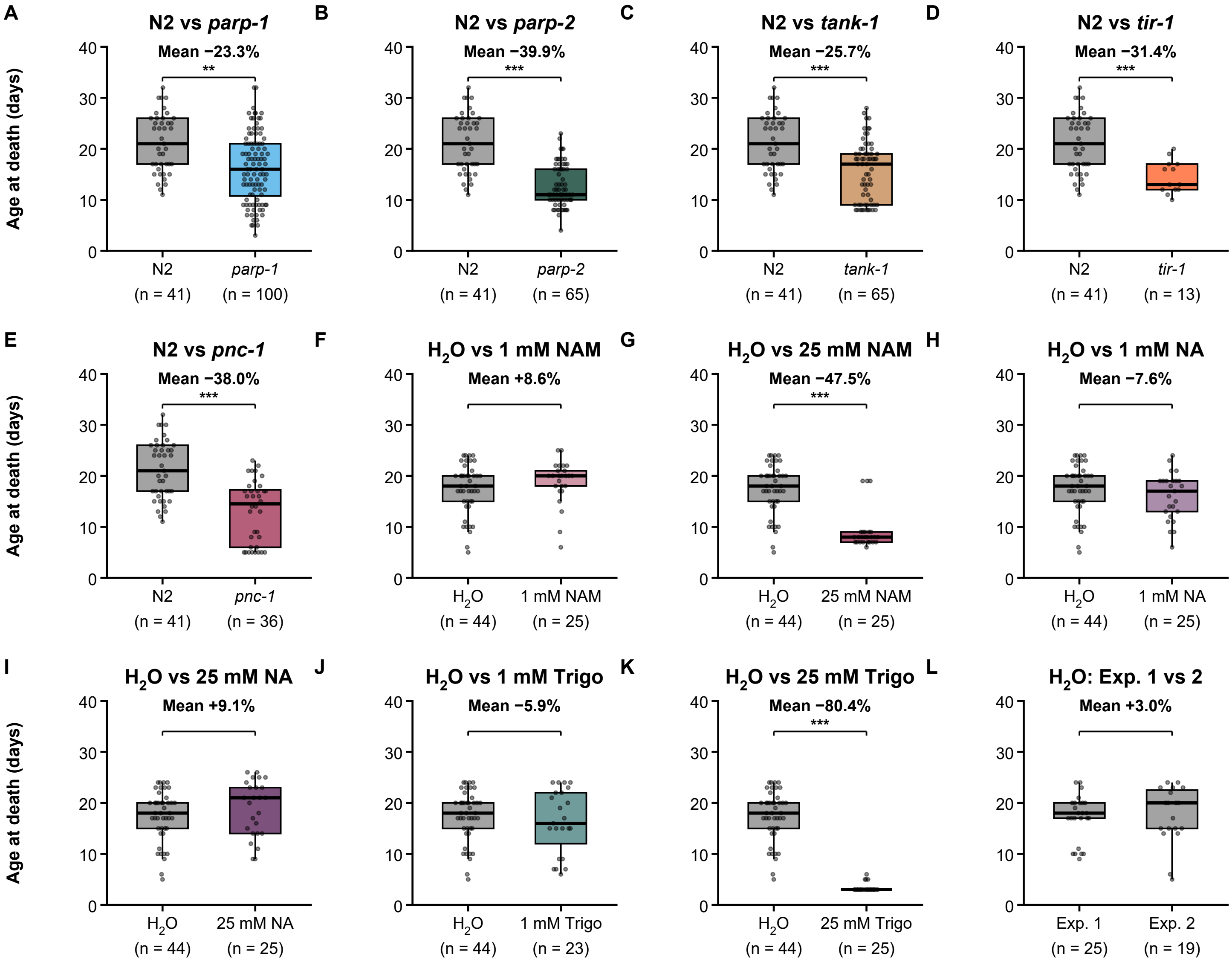

**Figure S7| Mean lifespan of $NAD^+$ pathway mutants and supplemented animals.**

Age at death of wild-type animals compared with *parp-1* (**S7A**), *parp-2* (**S7B**), *tank-1* (**S7C**), *tir-1* (**S7D**), and *pnc-1* (**S7E**) mutants, and of $H_2O$ vehicle controls compared with animals receiving 1 mM NAM (**S7F**), 25 mM NAM (**S7G**), 1 mM NA (**S7H**), 25 mM NA (**S7I**), 1 mM trigonelline (**S7J**), or 25 mM trigonelline (**S7K**). (**S7L**) Age at death of $H_2O$-treated controls from two independent experiments. Percentages indicate the change in mean lifespan relative to the control. Boxplots show medians and IQRs, whiskers show the full range, and points represent individual animals. Data are from the cohorts in Figure 7D and 7F; group sizes are shown below each group. Statistical significance was determined by pairwise log-rank (Mantel–Cox) tests with Holm correction. $^{**}p < 0.01$ and $^{***}p < 0.001$; unmarked comparisons were not significant.
